# Reprogramming enriches for somatic cell clones with small scale mutations in cancer-associated genes

**DOI:** 10.1101/2020.08.19.257238

**Authors:** Maike Kosanke, Katarzyna Osetek, Alexandra Haase, Lutz Wiehlmann, Colin Davenport, Adrian Schwarzer, Felix Adams, Axel Schambach, Sylvia Merkert, Stephanie Wunderlich, Sandra Menke, Marie Dorda, Ulrich Martin

## Abstract

Recent studies demonstrated that the mutational load in human induced pluripotent stem cells (hiPSCs) is largely derived from their parental cells, but it is still unknown whether reprogramming may enrich for individual mutations. 30 hiPSC lines were analyzed by whole exome sequencing. High accuracy amplicon sequencing showed that all analyzed small scale variants pre-existed in their parental cells and that individual mutations present in small subpopulations of parental cells become enriched among hiPSC clones during reprogramming. Among those, putatively actionable driver mutations affect genes related to cell cycle control, cell death and pluripotency, and may confer a selective advantage during reprogramming. Finally, a shRNA-based experimental approach was applied to provide additional evidence for the individual impact of such genes on the reprogramming efficiency. In conclusion, we show that enriched mutations in curated onco- and tumor suppressor genes may account for an increased tumor risk and impact the clinical value of patient-derived hiPSCs.

## Introduction

The availability of human induced pluripotent stem cells (iPSCs) (Takahashi et al., 2007) with their far-reaching potential for proliferation and differentiation offers novel opportunities for the development of tailored cellular therapies. Further research focusing on the genetic stability of reprogrammed cells is required, as considerable numbers of mutations in human iPSCs have been reported, and such genetic abnormalities might harbor the risk of tumor formation (Andrews et al., 2017; Weissbein et al., 2014). In general, small and large scale aberrations in iPSCs are thought to have at least three origins: i) mosaicism of pre-existing variants in the parental cell population, ii) mutagenicity of the reprogramming process itself, and iii) mutagenesis during prolonged culture.

Apparently, larger karyotype abnormalities and copy number variations arise in individual cells during culture expansion (Hussein et al., 2011; Ma et al., 2014; Mayshar et al., 2010), while individual ones can provide selective advantages and eventually dominate the population (reviewed by Andrews et al., 2017; Martin, 2017).

Besides larger aberrations, a high number of small genetic variants including single nucleotide variants (SNVs) and insertions and deletions (INDELs) have been detected in iPSCs (Bhutani et al., 2016; D’Antonio et al., 2018; Gore et al., 2011; Ji et al., 2012; Kwon et al., 2017; Lo Sardo et al., 2016; Young et al., 2012). Diverse experimental designs of recent studies, however, including the choice of somatic parental cell source, the passage number of analyzed iPSCs, the culture conditions, the type of abnormality examined and the analytic method led to contradictory results and a limited informative value of published data (Martin, 2017). Moreover, while next generation sequencing platforms differ in their error rates and ultimately their detection sensitivities (Goodwin et al., 2016), applied computational strategies develop fast and are far from being standardized, which further contributes to different conclusions concerning number and origin of variants in iPSCs (Goodwin et al., 2016; Lappalainen et al., 2019). Therefore, despite all efforts, selective clonal dynamics during reprogramming are poorly understood (Shakiba et al., 2019), and the extent, nature, and functional consequences of small genetic variants in iPSCs are still hardly characterized preventing any adequate assessment of risks associated with cellular therapies.

Several reports have hypothesized that the reprogramming process itself is ‘mutagenic’, and that small scale mutations are generated because of inefficient or inaccurate DNA repair mechanisms (Rouhani et al., 2016; Skamagki et al., 2017; Su et al., 2013), by replicative or by oxidative stress (Ben-David and Benvenisty, 2011; Ji et al., 2012; Rouhani et al., 2016; Sugiura et al., 2014; Yoshihara et al., 2017). Others reported that a substantial part or even the majority of small genetic variants in iPSC lines originate from their individual parental somatic cell clones. The proportion of genetic variants found to pre-exist in the parental cell cultures, however, varies among studies, since the sensitivities of the applied analytic methods differ. While older reports propose a high contribution of 50-74% of reprogramming-induced small scale variants to the mutational load of iPSCs (Gore et al., 2011; Ji et al., 2012; Rouhani et al., 2016; Sugiura et al., 2014), more recent studies report that around 90% of variants originate from somatic mosaicism in the parental cell population (D’Antonio et al., 2018; Kwon et al., 2017). It is still unknown, however, whether the remaining substantial number of undetectable genetic variants represent *de novo* mutations that have emerged during the reprogramming process (Rouhani et al., 2016; Yoshihara et al., 2017) or whether those ones might also pre-exist in the founder cell population as very rare variants not detectable by the applied techniques (Bhutani et al., 2016; Kwon et al., 2017).

The finding that the majority of small genetic variants in iPSCs originate from their parental somatic cell clones led to the speculation as to whether iPSCs derived from aged donors that accumulated *de novo* mutations over a lifetime (Vijg and Suh, 2013), may contain increased numbers of genetic aberrations. This speculation could not be confirmed by D’Antonio et al., who did not observe any correlation of donor age and mutational load in iPSCs (D’Antonio et al., 2018). In contrast, Lo Sardo et al. reported a cumulative number of variants with progressive donor age (Lo Sardo et al., 2016), and Skamagki demonstrated an elevated genomic instability in iPSCs derived from aged donors due reduction of DNA damage response by reactive oxygen species scavenge (Skamagki et al., 2017).

The parental origin of variants in iPSCs also raises the question whether specific mutations may provide a clonal selection advantage leading to enrichment of such mutations in iPSCs and to iPSC lines with altered cellular functions. In view of the various common characteristics of pluripotent stem cells and cancer (stem) cells, it can also be presumed that mutations, which provide a selection advantage during the reprogramming process, may lead to an increased tumor forming potential, in particular if cellular pathways are influenced that control cell cycling, apoptosis and pluripotency (Hadjimichael et al., 2015). Although the underlying molecular mechanisms remained largely unclear, this presumption was recently supported by a study of Shakiba et al., who observed that an ‘elite’ subset of dominating mouse embryonic fibroblast-derived clones overtook the whole cell population during reprogramming (Shakiba et al., 2019). With that finding, the study challenged the concept of clonal equipotency, where all clones have the same potential to attain iPSC state, and suggests that genetically encoded inequalities in cell fitness lead to dominance of otherwise hidden cells in the reprogramming niche (Shakiba et al., 2019). While Merkle et al. demonstrated a selective advantage of certain small scale mutations in the p53 gene during culture expansion of pluripotent stem cells (Merkle et al., 2017), there is, however, so far no evidence for selection of small genetic variants in any other gene during culture expansion, and recent studies could not demonstrate enrichment of any genetic variants from the parental cells during the reprogramming process (Cheng et al., 2012; Kwon et al., 2017; Lo Sardo et al., 2016; Rouhani et al., 2016; Ruiz et al., 2013; Young et al., 2012).

For a more adequate investigation of risks associated with clinical application of iPSC derivatives, we have generated a series of 30 early passage iPSC clonal lines from neonatal and aged individuals under controlled and comparable conditions to allow a systematic analysis for small scale variants via whole exome sequencing (WES). Importantly, we applied a sequencing technology with higher accuracy than the systems used in previous studies, and an ultra-sensitive amplicon sequencing approach to clarify the origin of detected variants and their frequency in parental cells. The most important aim of our study was to analyze whether individual variants and cell clones that carry such mutations in specific genes are enriched during the reprogramming process, and to what extent such variants are predicted to affect genes critical for cell function or cancer formation, which would call into question the general therapeutic usefulness of iPSCs (Ma et al., 2014).

## Results and Discussion

### Study design and characterization of small genetic variants in iPSCs

iPSC expansion-related enrichment of mutations (Merkle et al., 2017) was not in the focus of our study, we therefore restricted our analyses to early passage (mean P8-9, range P7-P12) clonal iPSC lines and their corresponding parental endothelial cell (EC) cultures. Overall, 30 iPSC clones (3 clones per donor to investigate intra-clonal variabilities) were analyzed (Fig. 1 and Table 1). Among them were 27 clones from umbilical vein ECs (hUVECs) or cord blood-derived ECs (hCBECs) of 6 neonatal donors, and from saphenous vein ECs (hSVECs) of 3 aged donors. In case of aged donor D#37, 3 additional iPSC clones derived from peripheral blood derived ECs (hPBECs) were analyzed (Table 1). For simplicity’s sake, hereinafter we will refer to D#37 hSVEC and D#37 hPBEC as individual ‘donors’. Direct whole exome sequencing (WES) was also performed for parental cells of two neonatal and two aged donors at the same passage as subjected to reprogramming (P4).

**Fig. 1:**
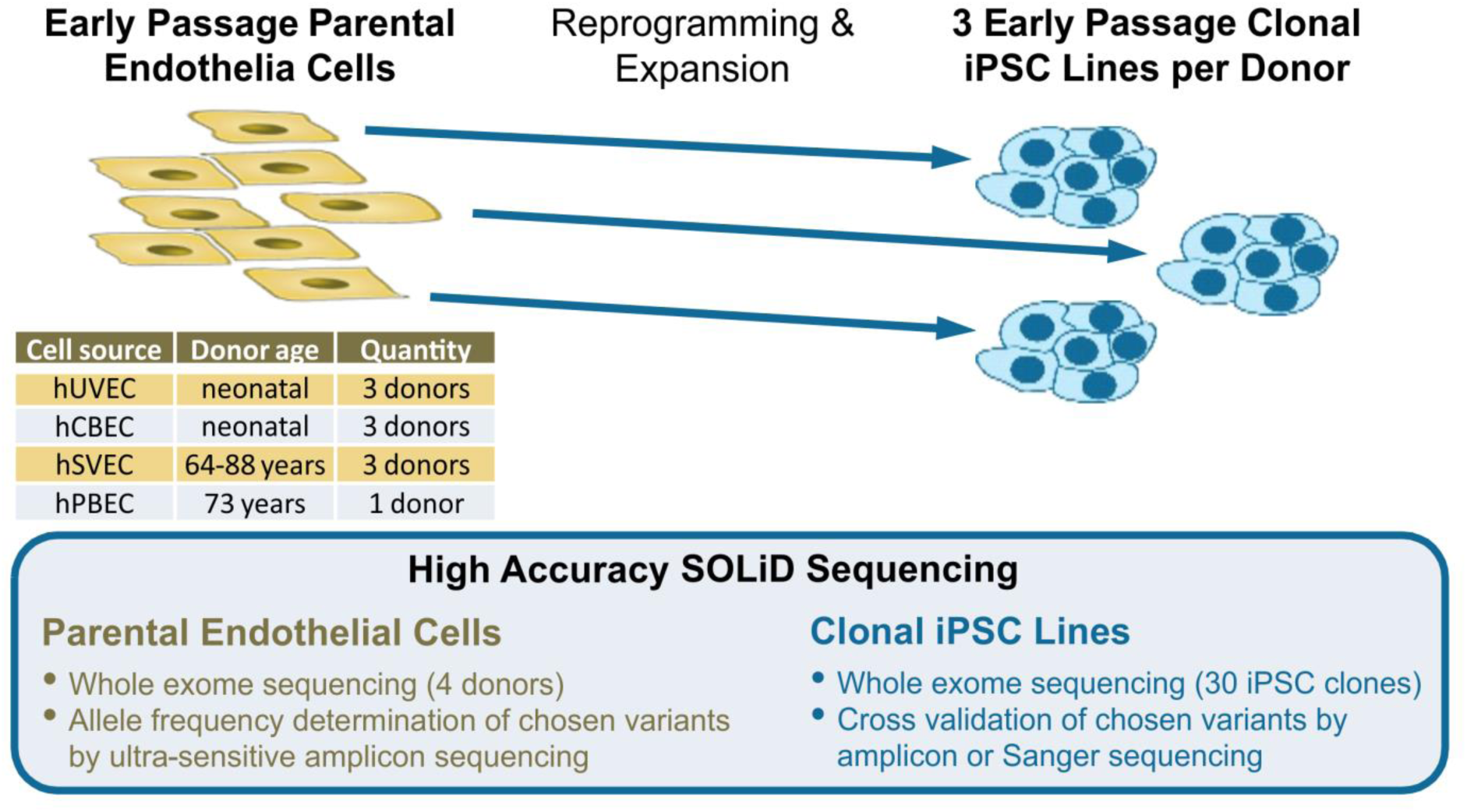
Study Design. 30 iPSC clones were generated from early passage endothelial cells (ECs), derived from, in total, 6 neonatal (umbilical vein, hUVEC; cord blood, hCBEC) and 3 aged donors (64-88 years, saphenous vein, hSVEC; peripheral blood, hPBEC). iPSC clones in passage 7-12 were subjected to whole exome sequencing (WES). 3 single cell iPSC clones per donor (in case of donor D#37, 3 clones derived from hSVECs and 3 clones from hPBECs) were included to investigate intra-clonal variabilities. Moreover, for 4 donors, variant frequency in the corresponding parental cell population was determined by WES to discriminate between enrichment of pre-existing variants and *de novo* mutagenesis during reprogramming. Additionally, allele frequencies of selected variants within parental cell populations were assessed by ultra-sensitive amplicon sequencing.

**Table 1:** Summary and categorization of small genetic variants detected in iPSCs from neonatal and aged donors. Exomes from 30 early passage clonal iPSC lines of neonatal and aged donors were sequenced. **Categories of variants:** A) Polymorphisms described in any population of GnomAD and 1000 Genome with minor allele frequency (MAF) ≥ 0.01. B) potentially common variants not listed as polymorphisms that were found in more than one donor, and C) donor-specific variants present in iPSCs of one donor, only. Potentially gene affecting variants (GAVs) compromise all variants in substantial gene regions such as coding and non-coding transcript region, UTRs, and splice regions, after exclusion of intergenic and intron variants. Abbreviations: EC, endothelial cells; hUVEC, human umbilical vein ECs; hCBECs, human cord blood derived ECs; hSVECs, human saphenous vein ECs; hPBECs, human peripheral blood-derived ECs.

|  |  |  | Variants |  |  |  |  |
| --- | --- | --- | --- | --- | --- | --- | --- |
| Donor | iPSC clone | Passage | Per Donor |  |  | Per iPSC clone |  |
|  |  |  | Total number | Polymorphisms and common variants (Cat. A&B) (%) | Donor-specific variants (Cat. C) | Donor-specific variants (Cat. C) | Potentially gene affecting variants (Cat.C GAVs) |
| iPSCs - Neonatal Donors |  |  |  |  |  |  |  |
| D#1 hUVEC | C2 | P8 | 37106 | 98.7 | 481 | 471 | 249 |
|  | C5 | P9 |  |  |  | 475 | 252 |
|  | C6 | P12 |  |  |  | 470 | 247 |
| D#2 hUVEC | C2 | P7 | 36354 | 98.6 | 514 | 507 | 293 |
|  | C4 | P7 |  |  |  | 504 | 291 |
|  | C5 | P7 |  |  |  | 503 | 288 |
| D#3 hUVEC | C1 | P9 | 42815 | 98.2 | 762 | 754 | 368 |
|  | C2 | P10 |  |  |  | 749 | 361 |
|  | C3 | P9 |  |  |  | 751 | 366 |
| D#22 hCBEC | C15 | P9 | 41226 | 98.4 | 645 | 634 | 296 |
|  | C17 | P9 |  |  |  | 631 | 296 |
|  | C21 | P10 |  |  |  | 631 | 295 |
| D#23 hCBEC | C1 | P7 | 34512 | 98.4 | 539 | 533 | 302 |
|  | C5 | P7 |  |  |  | 530 | 299 |
|  | C6 | P9 |  |  |  | 531 | 301 |
| D#25 hCBEC | C11 | P8 | 37776 | 98.5 | 559 | 555 | 286 |
|  | C12 | P8 |  |  |  | 552 | 285 |
|  | C13 | P8 |  |  |  | 555 | 286 |
| Mean |  |  | 38298 | 98 | 583 | 574 | 298 |
| iPSCs - Aged Donors |  |  |  |  |  |  |  |
| D#31 hSVEC (64 years) | C1 | P10 | 35509 | 98.7 | 465 | 454 | 251 |
|  | C3 | P9 |  |  |  | 452 | 250 |
|  | C4 | P8 |  |  |  | 450 | 248 |
| D#37 hSVEC (73 years) | C4 | P9 | 40296 | 98.4 | 626 | 594 | 292 |
|  | C8 | P9 |  |  |  | 600 | 292 |
|  | C10 | P9 |  |  |  | 602 | 297 |
| D#38 hSVEC (88 years) | C5 | P7+8 | 43292 | 97.8 | 946 | 920 | 291 |
|  | C6 | P9+10 |  |  |  | 920 | 290 |
|  | C9 | P8 |  |  |  | 921 | 292 |
| D#37 hPBEC (73 years) | C4 | P8 | 39588 | 98.4 | 620 | 613 | 301 |
|  | C14 | P9 |  |  |  | 616 | 304 |
|  | C15 | P10 |  |  |  | 616 | 305 |
| Mean |  |  | 39671 | 98 | 664 | 647 | 284 |
| iPSCs - All Donors |  |  |  |  |  |  |  |
| Mean |  |  | 38847 | 98 | 616 | 603 | 292 |

Importantly, our genome analyses were specifically designed to overcome limitations of recent studies with respect to detection and quantification of rare genetic variants in very small subpopulations of parental cells, and to prove potential enrichment of such mutations in iPSCs. The necessity to quantify these rare events against a large background of non-mutated DNA sequences requires highly accurate amplification and sequencing techniques. In light of that requirement, the inaccuracy of the standard “Sequencing by Synthesis” NGS systems with error rates of ∼ 1% (Liu et al., 2012) is a major limitation for the detection of a specific mutation carried by only a few cells among thousands of non-mutated genomes. In contrast to all previous studies (Bhutani et al., 2016; Cheng et al., 2012; D’Antonio et al., 2018; Gore et al., 2011; Ji et al., 2012; Kwon et al., 2017; Li et al., 2015; Lo Sardo et al., 2016; Rouhani et al., 2016; Su et al., 2013; Sugiura et al., 2014; Young et al., 2012), we developed a study design based on a SOLiD 5500XL system with an Exact Chemistry Call (ECC) module. The SOLiD system is the only system to use “Sequencing by Ligation” technology and has a substantially lower error rate (SOLiD 5500XL with ECC module: lower than 0,01% (Liu et al., 2012; Massingham and Goldman, 2012); see also SOLiD 5500XL Manual). To further reduce the number of false-positive variants, all analyzed iPSC clones were sequenced twice, and only genomic variants, which were detected in both runs were included in the further analyses. Moreover, we built a variant refinement strategy based on orthogonal validation sequencing (amplicon and Sanger sequencing). Cross-validation of, in total, 133 variants (Table S1 and Table S2) confirmed that our approach allowed reliable discrimination between sequencing artifacts and true variants. Although a number of true INDELs did not pass our filter criteria due to low mapping quality, in general INDELs up to 19 bp could be undoubtedly detected with our approach. Fig. S1A depicts the whole workflow, while further details on variant calling and refinement strategy are described in Methods. On average, 38847 variants were detected in iPSCs per donor (Table 1). Direct WES of iPSCs resulted in three categories of detected genetic variants: A) Polymorphisms described in any population of GnomAD and 1000 Genome with minor allele frequency (MAF) ≥ 0.01, B) potentially common variants not listed as polymorphisms that were found in more than one donor, and C) donor-specific variants present in 1, 2 or 3 iPSC clones of the respective donor (Fig. 2A).

**Fig. 2:**
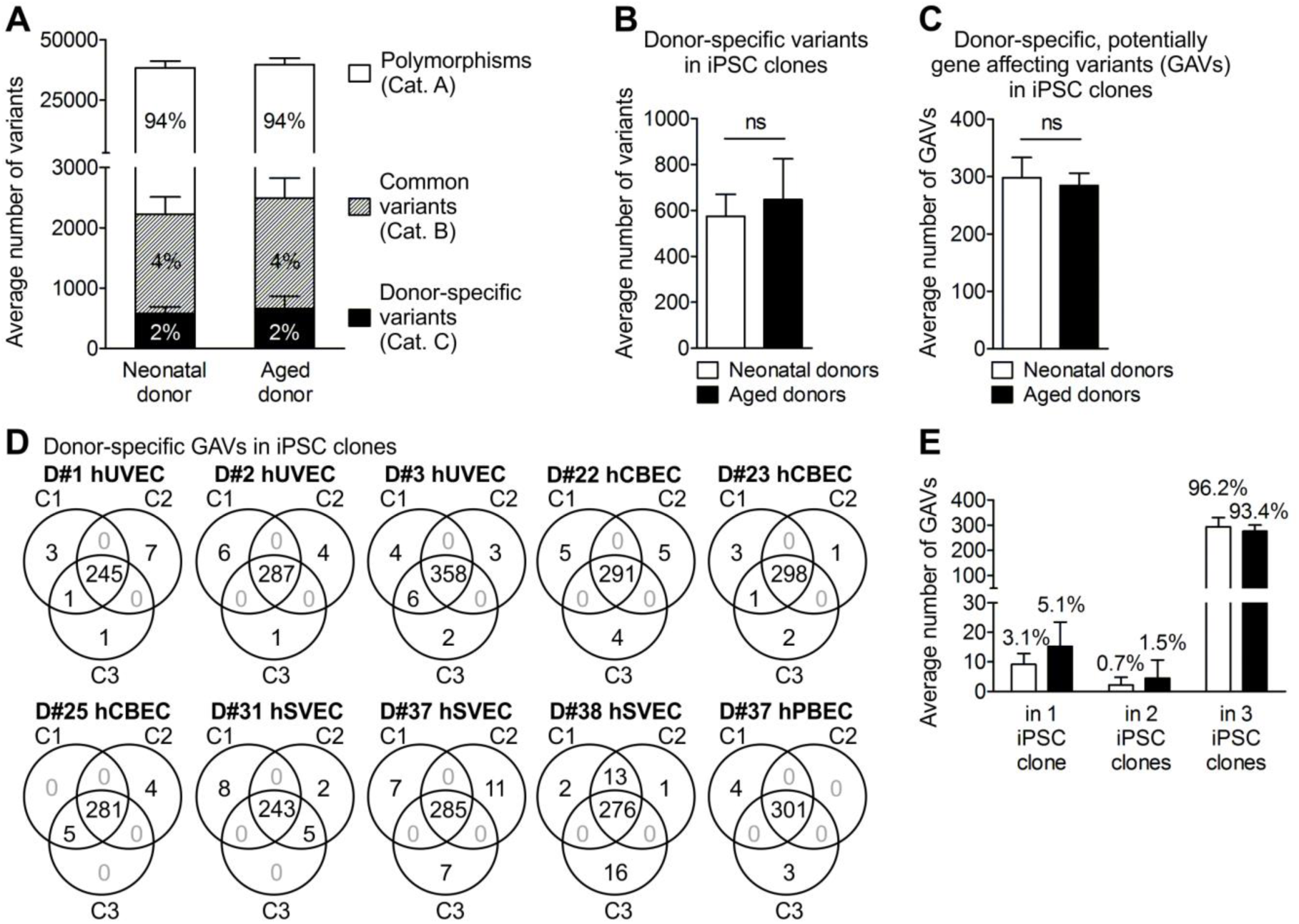
Distribution of small genetic variants in iPSC clones from neonatal and aged donors. **A** Averaged number of total genomic variants in 30 iPSC clones generated from neonatal and aged donors. For definition of variant categories see Table 1. **B** Averaged number of donor-specific variants (category C) per iPSC clone. Data were analyzed with unpaired two-tailed t-test; mean ± SD; n = 18 iPSC clones of neonatal donors, 12 of aged donors. **C** Averaged number of donor-specific, potentially gene affecting variants (GAVs, located in coding and non-coding transcripted regions, UTRs, and splice region) per iPSC clone derived of neonatal and aged donors. Data were analyzed with unpaired two-tailed t-test; mean ± SD; n = 18 iPSC clones of neonatal donors, 12 of aged donors. **D** Venn diagrams illustrating distribution and intersections of donor-specific GAVs between individual iPSC clones of the different donors. **E** Averaged dispersion of donor-specific GAVs in iPSC clones. Mean ± SD; n = 6 neonatal and 4 aged donors.

In agreement with previous reports (Gore et al., 2011; Young et al., 2012), about 98% of the detected genetic variants were common variants with 94% category A polymorphisms and 4% category B variants leaving 2% donor-specific category C variants (Table 1 and Fig. 2A, B). Despite enrichment for exomes, the pool of variants also included a substantial proportion of variants located in introns and intergenic regions. After elimination of such variants, on average, iPSC clones harbored 292 donor-specific, potentially gene affecting variants (GAVs) defined as variants located in the coding region, start or stop region, UTRs, splice regions, or non-coding transcript regions (Table 1 and Fig. 2C). While we cannot prove this for D#25 hCBEC C1 and C3, all other iPSC clones clearly represent independent, single cell derived clones as they all differ in terms of a number of unique GAVs (Fig. 2D, Fig. S2).

### iPSCs from aged donors do not contain significantly increased levels of total SNVs and INDELs

Lo Sardo et al. (Lo Sardo et al., 2016), who have generated iPSCs from *in vitro* expanded erythroid progenitors isolated from peripheral blood of adult donors of different age, recently reported increased numbers of small scale mutations in iPSCs derived from aged donors. While our experimental setting differed in the use of early passage ECs for reprogramming (Haase et al., 2009), we observed no significant difference in the number of variants in total (38298 vs 39671), donor-specific variants (574 vs 647) or GAVs (298 vs 284) per iPSC clone derived from neonatal or aged donors (Table 1 and Fig. 2A, B, C). A slight trend towards a higher number of donor-specific GAVs in iPSCs from aged donors was observed only for those detectable in 1 or 2 iPSC clones, only (Fig. 2E). These data are in general accordance with recent reports of D’Antonio et al. (D’Antonio et al., 2018). Interestingly, theoretical calculations demonstrated that the majority of cell divisions in humans already occur before birth (Frank, 2010): While there is an estimated number of 10^13^-10^14^ cells in a newborn (Bianconi et al., 2013; Frank, 2010) that require at least 45 cell divisions to be generated, 10^16^ cells are produced in total over the whole life time (Frank, 2010), which corresponds to an average of only ∼ 10 additional divisions per cell after birth. On top of this calculation, one has to consider that the reprogrammed ECs, both from neonatal and aged donors, underwent at least 10 sequential cell divisions between isolation from donor tissue and reprogramming of the expanded early passage cultures. Altogether, this may explain why we did not observe significant differences in the total number of mutations between neonatal and aged individuals.

Whether the contrary findings of Lo Sardo et al. (Lo Sardo et al., 2016) can be explained by the different sequencing technologies applied, or have their basis in the different parental cell types that have been reprogrammed, remains to be investigated. Actually, other studies indicated that the type of original somatic cell source and innate mosaicism may influence the mutational load of iPSCs (D’Antonio et al., 2018; Kwon et al., 2017; Su et al., 2013).

### Highly sensitive amplicon sequencing argues against appreciable contribution of *de novo* mutagenesis to small scale genetic variants detected in iPSCs

The majority of donor-specific variants in iPSCs (96.6%) including GAVs (95,1%) were detected in all 3 iPSC clones per donor representing donor-specific homo- or heterozygote variants (Fig. 2D, E and Table S3). A minority of variants, however, was detected in only 1 or 2 out of 3 iPSC clones from the respective donors (Fig. 2C, D and Table S3), a result that could be explained by three scenarios: i) genetic heterogeneity among the primary parental cell populations, ii) *de novo* mutation during reprogramming, and iii) reversal of heterozygous genetic variants e.g. by loss of heterozygosity (LoH) (Ryland et al., 2015) during reprogramming. Although the latter scenario cannot be entirely excluded, mitotic LoH is unlikely because it is a very rare event (Wijnhoven et al., 2001).

Direct WES of primary parental endothelial cell populations of 2 neonatal (D#2 hUVEC and D#3 hUVEC) and 2 aged (D#37 hSVEC and D#38 hSVEC) donors was performed in particular to exemplify the origin of donor-specific category C GAVs detected in iPSCs and to obtain evidence for the potential enrichment of rare variants from the parental cells. The sensitivity of the applied direct WES approach reached down to 10% of the diploid parental cell population (equals AF 0.05, Methods).

Overall, merely 7.1% of the donor-specific GAVs (80 SNVs and 5 INDELs, Fig. 3A and Table S4) could not be detected in the parental cell populations using this approach. Beside 7 GAVs, pre-existence of all variants detected in 3 out of 3 iPSC clones of the 4 donors in the founder cell population could be demonstrated by WES (Fig. 3B and Table S4). In contrast, it was not possible to statistically assure by WES the pre-existence in parental cells of any variant detected in 1 or 2 iPSC clones, only. According to these variants analyzed in parental cells from donors D#2 hUVEC, D#3 hUVEC, D#37 hSVEC and D#38 hSVEC, also all variants detected in 1 or 2 out of 3 iPSC clones of the remaining 6 donors including 64 GAVs were generally considered as undetectable in the parental cells by WES with frequencies below 10% (Table S5). The variants undetectable by WES could be either absent in the founder cell population, or present at allelic fractions below the sensitivity of our direct WES approach.

**Fig. 3:**
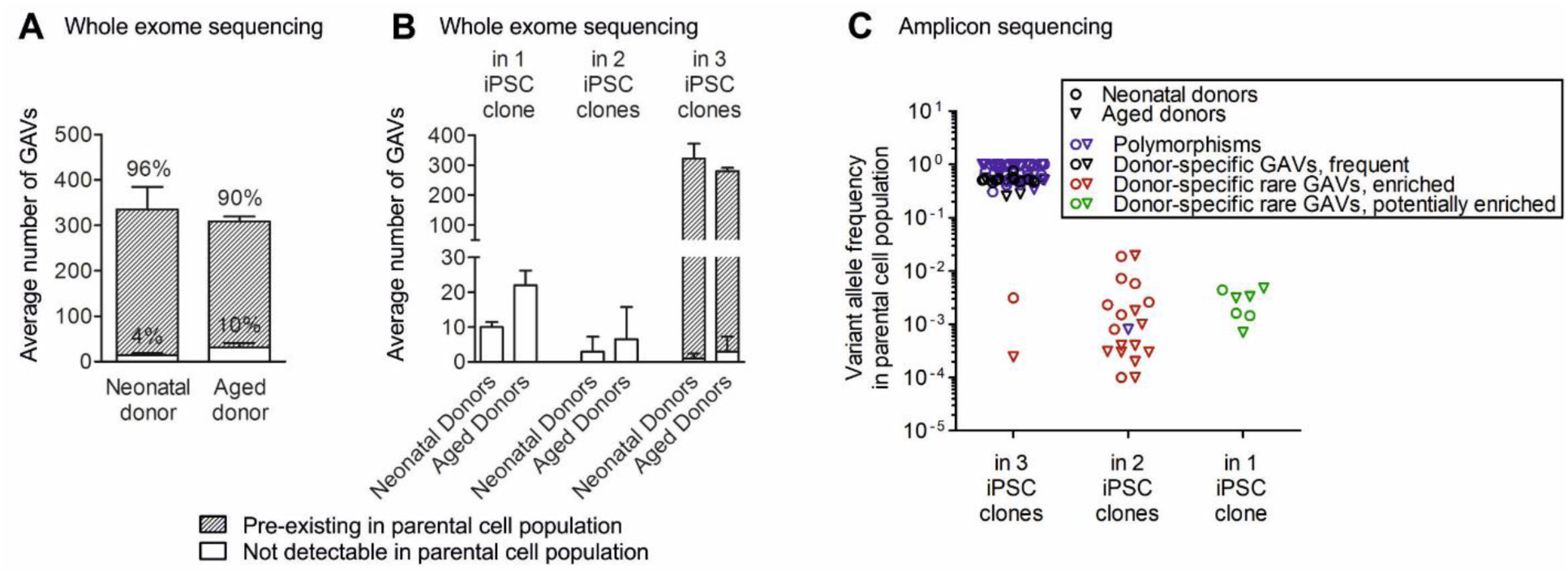
Reprogramming enriches for pre-existing genetic variants. **A** The origin of donor-specific, potentially gene affecting variants (GAVs) in iPSC clones was investigated in 4 donors by whole exome sequencing (WES) of the corresponding parental cell population. The majority of variants pre-existed in the parental cell population (detection limit of WES: 0.05 allele frequency). Mean ± SD; n = 2 neonatal and 2 aged donors. **B** Number of donor-specific GAVs present in 1, 2 or all 3 iPSC clones that are pre-existent or not detectable by WES in parental cells. Mean ± SD; n = 2 neonatal and 2 aged donors. **C** The pre-existence of 100 GAVs (45 polymorphisms, 6 common variants found in several donors, and 49 donor-specific variants) in the parental cell population was validated by amplicon sequencing (Table S2). The allele frequency of GAVs in the parental cell population is plotted against their presence in 1, 2 or 3 iPSC clones. Donor-specific GAVs were found shared between 2 or 3 iPSC clones of one donor, as well as being unique to one clone only. Donor-specific GAVs that were present in a small subpopulation in the respective parental cell population but were detected in 2 or 3 iPSC clones have evidently become enriched during reprogramming.

To assess their pre-existence in the corresponding parental cells, we selected a representative choice of 100 genomic variants detected by WES in iPSCs. 45 of these variants were listed in GnomAD or 1000 Genome as polymorphisms (category A) (Table S2) and 6 were not listed as polymorphisms but detectable in several donors (category B), 49 were also not listed as polymorphisms and were donor-specific (category C). Among the 49 donor-specific GAVs, 21 SNVs and 3 INDELs had been detected in 3 out of 3 iPSC clones, 16 SNVs and 2 INDEL in 2 out of 3 iPSC clones, and 7 SNVs and 0 INDELs in 1 out of 3 iPSC clones from the respective donor (Table S2). An optimized amplicon sequencing assay was established using the SOLiD 5500XL with ECC module. To obtain a realistic detection limit, we determined the overall error rate experimentally. Our amplicon sequencing approach exhibited an averaged coverage of 615274. Analysis of the non-mutated bases adjacent to the variant revealed an error rate of 0,12% and detection limit of 1 out of 2149 (SD 2564) reads to distinguish the existence of a SNV from average error (p value 0.1) (Table S2A). Application of this ultra-sensitive amplicon sequencing approach enabled us to confirm the pre-existence of all analyzed variants in the parental cell population, comprising 93 SNVs and 7 INDELs (Fig. 3C and Table S2). This finding strongly argues against appreciable *de novo* mutagenesis during reprogramming. In accordance with the reported high somatic mutation rates of 10^-7^ to 10^-6^ per gene per somatic cell division (Araten et al., 2005), our data indicate a substantial genetic heterogeneity with a multitude of rare SNVs and INDELs among the primary parental donor cell populations that are passed over to individual iPSC clones (Fig. 3).

Nevertheless, we cannot entirely negate the possibility of mutagenic effects during early passages of the reprogrammed cell clones. Since our exome sequencing analyses of the iPSC clones were limited to variants that exist at least in one clone with a frequency > 0.3, we would have excluded variants that might have developed after the single cell cloning step while the reprogramming process may not have been fully completed.

### Reprogramming enriches for individual variants located in cancer-associated genes

The vast majority of variants were detected in 3 out of 3 iPSC clones (categories A or B, and most of the donor-specific variants) and showed a high allele frequency in their parental cells, indicating homo- or heterozygosity in the majority of the respective cell population.

18 GAVs that had been detected in 2 of 3 iPSC clones of donors D#3 hUVEC, D#25 hCBEC D#31 hSVEC and D#38 hSVEC (13 of those also analyzed and undetected by WES of parental cells) were further analyzed by amplicon sequencing. Remarkably, all these GAVs were detectable in the parental cells at low frequency (0.02-4% of the diploid founder cell population, Table S2), strongly suggesting enrichment during the reprogramming process. It can be excluded that the presence of these 18 GAVs in 2 out of 3 iPSC clones simply represent a statistical phenomenon for two reasons: i) Without enrichment, the likelihood to find a rare variant with a frequency between 0.02 and 4% in 2 out of 3 iPSC clones of one donor is between P=4*10^-7^ and P=1.5*10^-2^ (on average P=2.6*10^-4^), only. ii) Supposed that appearance of these 18 GAVs in 2 out of 3 iPSC clones represent just a statistical phenomenon based on a very high number of rare variants in the parental cell populations, one would expect that a much higher number of such rare GAVs should be detectable in 1 out of 3 iPSC clones than in 2 out of 3 iPSC clones, which was not observed in our study (the likelihood P for presence in iPSC clone is on average 193 fold higher than for presence in 2 out of 3 iPSC clones). Therefore, the 18 GAVs mentioned above, as well as 2 GAVs that had been detected in 3 out of 3 iPSC clones derived from D#3 hUVEC and D#38 hSVEC and were confirmed to be rare in their parental cell population by amplicon sequencing (undetectable by WES) (Table S2) are termed afterwards ‘enriched GAV’. Since all analyzed 18 GAVs that were detected in 2 out of 3 iPSC clones proved to pre-exist only in a small subpopulation of parental cells, also all other 13 category C GAVs detected in 2 out of 3 iPSC clones were further considered generally as ‘enriched’ (Table S5A), leading to a total number of 31 ‘enriched GAVs’. Based on that considerations, iPSCs overall comprise between 0 and 18 enriched GAVs per clone (Table S5A).

In addition, 108 donor-specific category C GAVs were detected in 1 out of 3 iPSC clones, only, of the 10 donors (Table S5B). As far as analyzed, these GAVs were undetectable by direct WES but were shown be present with low frequency in the parental cells by amplicon sequencing. Although our approach did not allow to draw any direct conclusion about enrichment of individual GAVs of this group, it can be presumed that also a considerable proportion of these variants had been enriched to a certain extent during reprogramming.

Furthermore, we analyzed whether the transition/transversion (Ts/Tv) ratio and mutation spectra of donor-specific and especially enriched GAVs may point to an *in vitro* or *in vivo* origin of the respective variants. In fact, increased oxidative stress can result in C>A transversions (Viel et al., 2017) and is frequently observed during *in vitro* culture of somatic cells. An elevated frequency of C>A transversions is therefore considered as typical *in vitro* signature. Remarkably, we observed an overall increased proportion of C>A transversions among GAVs detected in 1 out of 3 iPSC clones (Fig. 4A), suggesting that the *in vitro* expansion culture of ECs prior to reprogramming contributed to the mutational load.

**Fig. 4:**
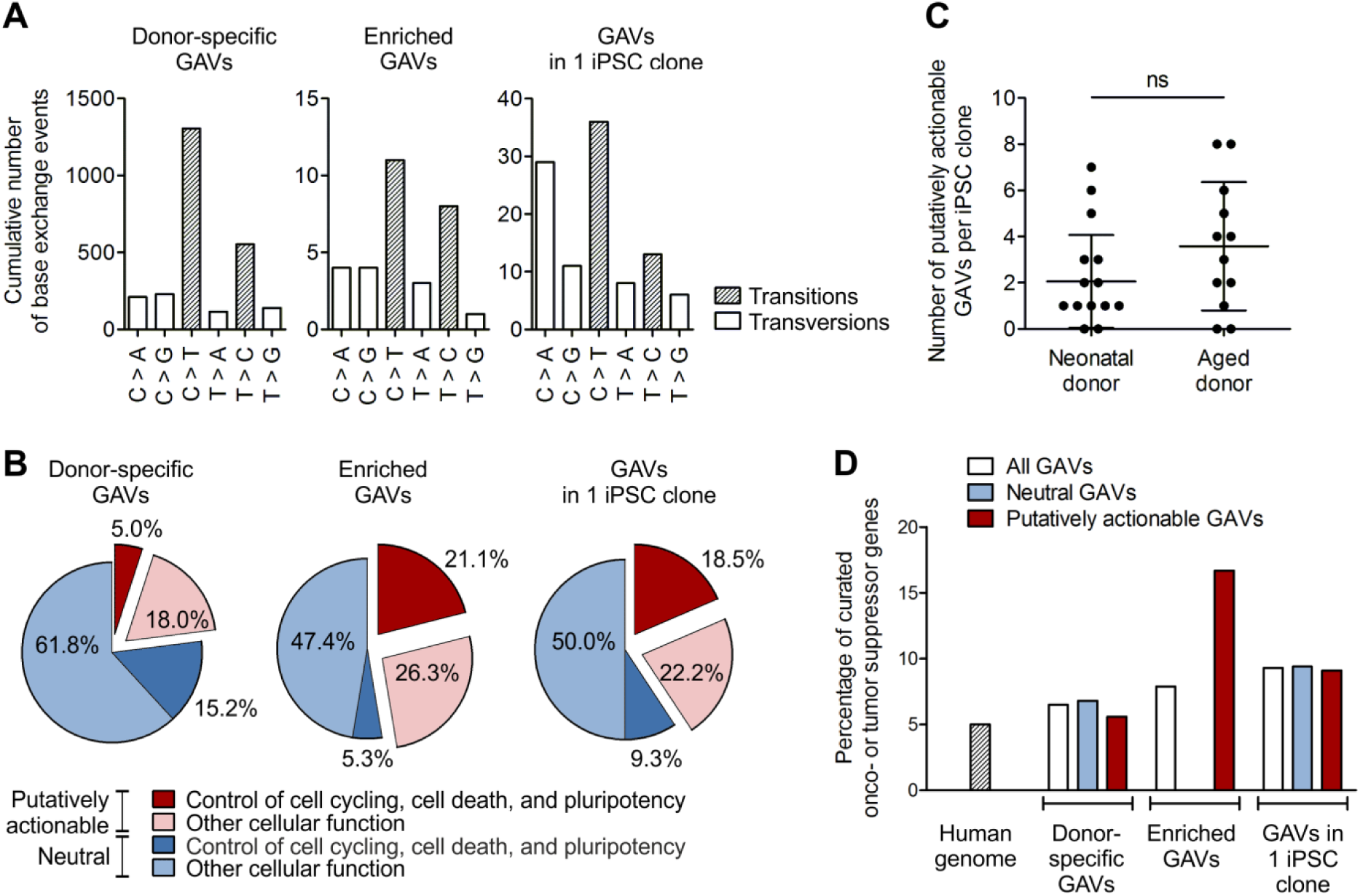
Reprogramming enriches for putatively actionable mutations in genes connected to cell cycling, cell death or pluripotency, and in curated onco- or tumor suppressor genes. The nature of donor-specific category C, potentially gene affecting variants (GVAs) detected in iPSC clones was characterized. **A** Mutational spectra of the collective of donor-specific GAVs, enriched GAVs and GAVs unique to 1 iPSC clone. n = 10 donors. **B** Percentage of putatively actionable GAVs, defined by harmful designation predicted by consensus of in silico prediction of Condel, FATHMM, CADD and SnpEff impact (red) as well as proportion of neutral GAVs (blue). The percentage of variants that affect genes involved in control of cell cycling, cell death, and pluripotency is depicted in dark red or blue, respectively. The proportion of variants in genes with other cellular functions is plotted in light red or blue. n = 10 donors. **C** Total number of putative actionable enriched GAVs, and GAVs unique to only 1 iPSC clone, per iPSC clone. Mean ± SD; n = 18 iPSC clones of neonatal donors, 12 of aged donors; discrepancy of samples was assessed applying two-tailed Mann Whitney test. **D** Percentage of curated onco- or tumor suppressor genes as found in the human genome (calculated based on COSMIC cancer gene census and OncoKB database), or affected by neutral or putatively actionable, donor-specific GAVs, enriched GAVs, or GAVs in 1 iPSC clone.

We also explored whether the mutation spectra of variants and especially enriched GAVs may correlate with specific mutational signatures of cancer. Actually, different mutational processes that include DNA damage and inaccurate maintenance mechanisms are considered to act with variable strength throughout the lineage specification and evolution of cancer cells (Alexandrov and Stratton, 2014; Ju et al., 2017). The overall Ts/Tv ratio of total variants, donor-specific variants, and donor-specific GAVs in the iPSC clones was 2.5, 2.2, and 2.7, respectively, which is consistent with the reported ratios of 2-2.1 for whole genome and 3.0 for human exonic regions (Marth et al., 2011; Zhang et al., 2015). In agreement with the results of Kwon et al. (Kwon et al., 2017), the entirety of donor specific category C GAVs found in all 3 iPSC clones were dominated by C>T transition without strand-bias (Fig. 4A). Such C>T transitions as principal nucleotide changes indicate spontaneous deamination of 5-methylcytosine and are a hallmark of signature 1 of “Signatures of Mutational Processes in Human Cancer” (COSMIC Catalogue of somatic mutations in cancer), which is the result of an endogenous mutational process present in most normal (and neoplastic) somatic cells. Interestingly, enriched GAVs were characterized by an increased T>C transition without strand-bias pointing to mutational signatures 6, 15, 20 and 26, which are often found in different cancer types and are all caused by defective DNA mismatch repair during replication of somatic cells (COSMIC Catalogue of somatic mutations in cancer), supporting the parental cell origin of these mutations (Fig. 4A).

Finally we analyzed the entirety of donor-specific (category C) GAVs and all enriched GAVs for functional consequences on the affected genes. In general, it can be expected that enrichment of certain genetic variants in iPSCs implies that individual variants affect genes, pathways and cellular functions that influence the reprogramming process. However, the observation that the analyzed iPSCs contain a substantial number of, on average, 2.8 (range 0-18) of enriched variants per clone suggests that only some of those variants actually operate as driver mutations, while others are passenger variants. Indeed, prediction of variant impact on gene functionality based on a consensus of Condel, FATHMM, CADD, and SnpEff in silico prediction tools revealed 53% neutral variants among the enriched GAVs, which most likely constitute passenger variants, and 47% putatively actionable enriched GAVs (Fig. 4B and Table S5A). Remarkably, this proportion of putatively actionable GAVs among the enriched GAVs is much higher than within the entirety of category C GAVs found in 3 of 3 iPSC clones (23%), further supporting the hypothesized active role of putatively actionable GAVs in the enrichment process during reprogramming. While 18 putatively actionable enriched GAVs were detected in 4 of the donors, only (Table 2), D#38 hSVEC for instance harbors even 7 putatively actionable enriched GAVs including mutations in the curated onco- and tumor suppressor genes JAK1 and XIAP (COSMIC Cancer gene census and OncoKB). Both mutations of the genes JAK1 and XIAP affect substantial protein domains thereof: The mutation in catalytic phosphotransferase domain (Uniprot, Pfam) of tyrosine-protein kinase JAK1 might affect diverse signaling pathways including interleukin, EGFR1, type II interferon, and TGF beta signaling pathways and alter cellular processes including proliferation, differentiation, or apoptosis (COSMIC, Reactome, WikiPathway). As onco- and tumor-suppressor gene JAK1 is reported to mediate “Escaping programmed cell death” and “Proliferative signalling” (COSMIC Hallmarks of Cancer). Similarly, the E3 ubiquitin-protein ligase XIAP harbors a putatively actionable mutation within its BIR1 domain, which is crucial for homodimerization with TAB1, and recruitment of TAK1, an important regulatory component of the NF-κB canonical pathway promoting cell survival (Lu et al., 2007; Sorrentino et al., 2019). Moreover, mutation in the BIR1 domain (Pfam) might reduce SMAC binding, caspase release from XIAP, and induction of cell death (Attaran-Bandarabadi et al., 2017; Lu et al., 2007). We therefore presume that these putatively actionable GAVs confer a selective advantage and drive the reprogramming of cells carrying such mutations, whereas the neutral GAVs more likely represent passenger mutations coincidentally coexisting in the somatic parental cells.

**Table 2:** Reprogramming enriches for pre-existing genetic variants in curated onco- and tumor suppressor genes, and in genes connected to cell cycling, cell death or pluripotency. The table lists all putatively actionable GAVs found in 3 or 2 iPSC clones but not detected in parental cell population by WES (upper part), or found in 2 iPSC clones but were not analyzed in parental cell population via WES (lower part) that have been defined as enriched variants. Ultra-sensitive amplicon sequencing of GAV spanning regions for precise determination of allelic frequencies in the corresponding parental cell population was performed for a representative choice of variants. Pre-existence of variants in parental cell population was confirmed (p-value 0.1) taking local error rates into account (Table S2A). Functional consequence of potentially gene affecting variants (GVAs), was classified by a consensus of the in silico prediction of Condel, FATHMM, CADD and SnpEff impact. Curated onco- or tumor suppressor genes (annotation retrieved from OncoKB or COSMIC Cancer gene census) are shaded in red. Variants that are presumed to influence reprogramming since they affect genes with function in control of cell cycling, cell death or pluripotency are depicted in blue font. () Pre-existence is very likely but not confirmed with statistical confidence.

| Gene symbol | Chromosome | Start position | Reference genotype | Variant genotype | Donor | iPSC clone | Allele frequency in iPSC clones by exome sequencing | Allele frequency in parental cells by exome sequencing | Allele frequency in parental cells by amplicon sequencing | Consequence / Variant type | GO Processes | Mutations in affected genes previously detected in pluripotent stem cells |
| --- | --- | --- | --- | --- | --- | --- | --- | --- | --- | --- | --- | --- |
| GOLGB1 | 3 | 121433800 | C | T | D#3 hUVEC | C1<br>C2<br>C3 | 0.41<br>0.33<br>0.43 | 0 | 0.0031 | Missense | endomembrane system organization / establishment of localization in cell |  |
| UACA | 15 | 70959677 | G | T | D#3 hUVEC | C1<br>C2<br>C3 | 0.35<br>0.36<br>0.36 | 0 | nd | Missense | cell death |  |
| KIF1A | 2 | 241712557 | G | C | D#3 hUVEC | C1<br>C3 | 0.42<br>0.64 | 0 | 0.0015 | Missense | establishment of localization in cell / microtubule based process | Lo Sardo, Nat Biotechnol. (2017) |
| HYOU1 | 11 | 118923042 | CA | C | D#3 hUVEC | C1<br>C3 | 0.36<br>0.28 | 0 | 0.0026 | Frameshift | response to stress / establishment of localization in cell / cell death |  |
| SLCO1B7 | 12 | 21200041 | TA | T | D#3 hUVEC | C1<br>C3 | 0.39<br>0.35 | 0 | 0.0001 | Frameshift | anion transport | lhry, Cell Rep. (2019) |
| SLC8A3 | 14 | 70634061 | C | T | D#3 hUVEC | C1<br>C3 | 0.39<br>0.44 | 0 | 0.0023 | Missense | homeostasis / transmembrane transport / response to stress / establishment of localization in cell / ion transport |  |
| PPP1CC | 12 | 111160453 | G | T | D#38 hSVEC | C5<br>C6<br>C9 | 0.29<br>0.33<br>0.69 | 0 | 0.0002 | Missense / stop gain | cell cycle / cell division / chromosome segregation / metabolic process / regulation of transport | lhry, Cell Rep. (2019) |
| JAK1 | 1 | 65300295 | G | C | D#38 hSVEC | C5<br>C6 | 0.33<br>0.48 | 0 | 0.0003 | Missense | catalytic activity / cell proliferation / response to stress / response to cytokine / signal transduction |  |
| CEP350 | 1 | 180062780 | C | T | D#38 hSVEC | C5<br>C6 | 0.39<br>0.48 | 0 | 0.0003 | Stop gained | cytoskeleton organization / microtubule anchoring / microtubule based process |  |
| MYO7A | 11 | 76924004 | C | T | D#38 hSVEC | C5<br>C6 | 0.51<br>0.47 | 0 | 0.0004 | Missense | cell morphogenesis involved in differentiation / establishment of localization in cell / protein transport / organelle assembly / phagocytosis | Lo Sardo, Nat Biotechnol. (2017); Rohani, PLoS Genet. (2016) |
| SCEL | 13 | 78184187 | A | G | D#38 hSVEC | C5<br>C6 | 0.38<br>0.32 | 0 | 0.0010 | Missense; 3' UTR | tissue development |  |
| HIRIP3 | 16 | 30004693 | T | A | D#38 hSVEC | C5<br>C6 | 0.10<br>0.33 | 0 | nd | Splice site (acceptor) | chromatin assembly or disassembly / chromosome organization |  |
| XIAP | X | 123019673 | A | G | D#38 hSVEC | C5<br>C6 | 1.00<br>0.98 | 0 | 0.0003 | Missense | response to DNA damage stimulus / response to stress / cell cycle / spindle assembly / cell death / regulation of gene expression / response to growth factor stimulus / cell proliferation | Kilpine, Nature (2017) |
| ZBED9 | 6 | 28539848 | G | T | D#25 hCBEC | C11<br>C13 | 0.39<br>0.41 | nd | nd | Stop gained | DNA integration |  |
| TRIM9 | 14 | 51477193 | G | T | D#25 hCBEC | C11<br>C13 | 0.54<br>0.44 | nd | 0.0187 | Missense | catabolic process / cell cell signaling / establishment of localization in cell / exocytosis / regulation of transport / proteolysis / vesicle mediated transport |  |
| MMP9 | 20 | 44642429 | T | C | D#25 hCBEC | C11<br>C13 | 0.57<br>0.36 | nd | 0.0058 | Missense | catabolic process / cell death / signal transduction / ion transport / regulation of dna binding / mitochondrion organization / release of cytochrome c from mitochondria |  |
| CSMD3 | 8 | 113308154 | T | C | D#31 hSVEC | C3<br>C4 | 0.48<br>0.50 | nd | 0.0018 | Missense | Interact with NEK4 (cell division / response to DNA damage stimulus / cell cycle / regulation of cellular senescence / regulation of gene expression) | Lo Sardo, Nat Biotechnol. (2017) |
| SALL1 | 16 | 51172618 | G | A | D#31 hSVEC | C3<br>C4 | 0.48<br>0.54 | nd | 0.0194 | Missense | chromatin modification / chromosome organization / growth / maintenance of cell number / regulation of gene expression / biosynthetic process / response to stimulus / somatic stem cell population maintenance | lhry, Cell Rep. (2019); Bhutani, Nat Commun. (2016); Gore, Nature. (2011) |

Furthermore, we determined the proportion of GAVs that affect genes involved in control of cell cycling, cell death and pluripotency networks (GO process annotation based classification), which can be expected to impact reprogramming efficiency. According to GO annotation ∼ 20.8% of all human genes belong to this group (calculated based annotation in GO biological process (C5BP) collection of MSigDB C5). The observed proportion of 20.2% (15,2 + 5,0%) of such GAVs among the entirety of category C GAVs correspond very well to that (Fig. 4B left graph). Remarkably, these genes are considerably overrepresented among the enriched GAVs with 26.3% (21,1 + 5,3%; Fig. 4B middle graph). Even more striking is their contribution within the putatively actionable enriched GAVs: here, 44.4% of variants (equals to 21.1% of all enriched GAVs) affect genes involved in control of cell cycling, cell death and pluripotency networks, while only 10,1% enriched neutral GAVs (equals to 5.3% of all enriched GAVs; Fig. 4B middle graph) affect such genes. In contrast, the entirety of putatively actionable category C GAVs contained only 21.6% of such genes (equals to merely 5% of all donor-specific GAVs) (Fig. 4B left graph).

Besides obviously enriched GAVs, we identified 44 putatively actionable GAVs unique to 1 out of 3 iPSC clones (Table 5B) which accounts for 40.7% (18.5% + 22.2%) of all GAVs found in 1 clone (Fig. 4B right graph). Remarkably, also in this group a disproportionately high number of genes involved in control of cell cycling, cell death and pluripotency networks were identified (45,5%) (equals to 18.5% of all GAVs unique to 1 iPSC clone), which is in contrast to their neutral counterpart (15,6%) (equals to 9.3% of all GAVs unique to 1 iPSC clone; Fig. 4B, right graph), supporting the presumption that also many of the GAVs unique to 1 out of 3 iPSC clones had been enriched during reprogramming.

Altogether, ∼ 87% of iPSC clones harbored 1-7 putatively actionable GAVs, in ∼ 67% of iPSC clones located in genes with pivotal function for cell cycling, cell death, or pluripotency (Table S5). Interestingly, in contrast to the overall number of donor-specific (category C) variants and GAVs, iPSC clones from aged donors harbored a higher proportion of putatively actionable GAVs in genes involved in cell cycling, cell death or pluripotency (75%) than clones derived from neonatal donors (61%). Moreover, iPSCs derived from aged donors exhibited, on average, slightly more putatively actionable, biologically relevant mutations per clone (mean 2.1 and median 1 compared to 3.6 and 3.5 in iPSC clones of aged donors) (Fig. 4C).

In addition, we analyzed the proportion of variants in our iPSCs affecting oncogenes (OGs) and tumor suppressor genes (TSGs). According to OncoKB or COSMIC Cancer gene census database, the human genome contains 942 curated OGs / TSGs representing ∼ 5% of all human genes. A very similar proportion of variants affecting OGs / TSGs (6.5%) were observed in the entirety of donor-specific GAVs detected in all 3 clones (Fig. 4D). Enriched GAVs, however, contained a considerably higher percentage (17%) of putatively actionable GAVs located in curated OGs / TSGs, namely JAK1, XIAP, CSMD3 (Fig. 4D, Table 2, Table S5A shaded in red), suggesting that reprogramming also enriches for specific mutations in oncogenes.

Similar to the increased proportion of GAVs affecting genes involved in control of cell cycling, cell death and pluripotency, GAVs unique to 1 iPSC clone also contained an increased percentage (9%) of variants in curated OGs / TSGs (4 variants in GRIN2A, TCF12, XPC, and SOX2, Fig. 4D, Table S5B shaded in red).

Interestingly, some of the enriched mutations and potentially enriched mutations (detected in 1 out of 3 iPSC clones) that have been observed in our study affect genes that are also reported by previous studies to be mutated in iPSCs, although their relevance was, in general, not further recognized. In those studies the respective variants had been considered as derived from a small subpopulation of the source cells during reprogramming, or, if not detectable in the parental cells, *de novo* mutations (Bhutani et al., 2016; Gore et al., 2011; Ishikawa, 2017; Ji et al., 2012; Kilpinen et al., 2017; Kwon et al., 2017; Lo Sardo et al., 2016; Rouhani et al., 2016; Young et al., 2012), or were reported to affected iPSC fitness (Ihry et al., 2019). The recurrently affected genes comprise KIF1A (Lo Sardo et al., 2016), SLCO1B7 (Ihry et al., 2019), MYO7A (Lo Sardo et al., 2016; Rouhani et al., 2016), XIAP (Kilpinen et al., 2017), CSMD3 (Lo Sardo et al., 2016), SALL1 (Bhutani et al., 2016; Gore et al., 2011; Ihry et al., 2019), all affected by enriched mutations in our work, and GRIN2A (Lo Sardo et al., 2016; Rouhani et al., 2016), CCDC180 (Ihry et al., 2019), TCF12 (Gore et al., 2011), ATF7IP (Rouhani et al., 2016), HDAC3 (Ihry et al., 2019), PPRC1 (Ihry et al., 2019), TTN (Ishikawa, 2017; Ji et al., 2012; Young et al., 2012), SOX2 (Kwon et al., 2017), ABCB11 (Lo Sardo et al., 2016), and PDE8A (Lo Sardo et al., 2016), which also carried mutations unique to 1 out of 3 of our iPSC clones. Some of those recurrently mutated genes represent curated OGs / TSGs (XIAP, CSMD3, GRIN2A, TCF12 and SOX2). Further genes, such as SOX2 or SALL1, are involved in the molecular control of pluripotency (Ractome.org; R-HSA-2972975.1: POU5F1 (OCT4), SOX2, NANOG Bind the SALL1 Promoter) or cell cycling (CCDC180, HDAC3, TTN; Table S5B). The predicted functional impact of these mutations and the biological relevance of affected genes suggest selective advantages as mechanism for enrichment of somatic cell clones carrying such a mutation during reprogramming.

Lastly, we aimed to experimentally confirm the impact of such genes, which were affected by enriched putatively actionable GAVs, on the reprogramming process. Among those genes (Tab. 2), in particular SALL1, a tumor suppressor (Ma et al., 2018), appeared of interest because mutations in SALL1 were also recently observed in hiPSCs by others (Bhutani et al., 2016; Gore et al., 2011) or as pluripotency-specific gene (Ihry et al.). Furthermore, SALL1 interacts with OCT4 (Pardo et al., 2010), SOX2 (Karantzali et al., 2011) and NANOG (Lopes Novo and Rugg-Gunn, 2016) as key factors of reprogramming (Ihry et al., 2019). Hence, we investigated the effect of a knockdown of SALL1 and 15 other genes on the reprogramming efficiency by utilizing a lentiviral vector-based RNA interference system that delivers short hairpin RNAs embedded in an optimized human miR-30 backbone (shRNAmiRs) to achieve stable and heritable sequence-specific gene knockdown (Adams et al., 2017). For this purpose, one of our parental cell populations, D#2 hUVECs, was transduced with a library containing 48 shRNAmiRs against those 16 genes (3 shRNAmiRs / gene). Besides SALL1, those 16 genes include ZFHX3, a curated oncogene, which is also directly interacting with OCT4 (Pardo et al., 2010) and was chosen as internal positive control, and a representative choice of genes that were identified in our iPSC clones to be affected by Cat. A (ALMS1, ACTR8), Cat. B (CCDC14, KCTD8, KLHL14) (Table S1) or enriched Cat. C variants (putatively actionable: CEP350, SLCO1B7, MAGEB6, MYO7A and SCEL, neutral or unconfirmed: SLC12A4, TMEM 139, TMEM168, TNNI3K) (Table S6). Next, 3 batches of the transduced cells were independently reprogrammed and the composition of shRNAmiRs in the cell population before and after reprogramming (P3) was quantified. While in the transduced parental cell population the distribution of all shRNAmiRs was homogenous (mean frequency 0.019, SD 0.008), in the 3 iPSC batches only a few shRNAmiRs dominated the population (Fig. 5A). These shRNAmiRs were found > 2 fold enriched compared to their frequencies within the parental cell population (Fig. 5B, red bar) in 1 or even 2 iPSC batches. In contrast, the frequency of most shRNAmiRs (77%) remained on a similar level or was reduced in the iPSC batches (fold change < 2) (Fig. 5B, black bars).

**Fig. 5:**
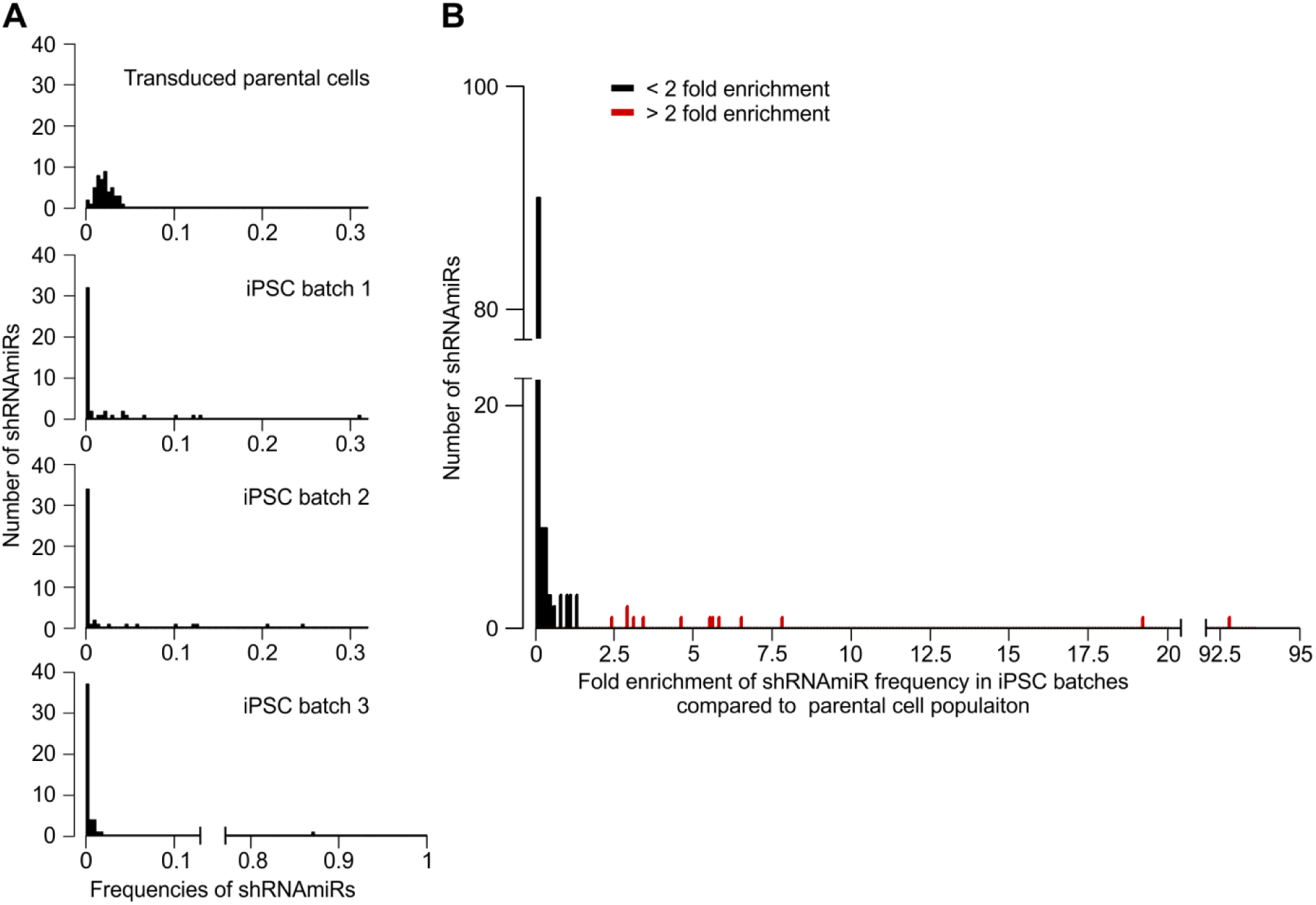
Reprogramming leads to enrichment of individual shRNAmiRs directed against genes that were affected by enriched, putatively actionable GAVs. For a multiplexed shRNAmiR screen, D #2 hUVECs were transduced with a library containing shRNAmiRs against a choice of 16 genes (3 shRNAmiRs / gene) before cells were reprogrammed in 3 independent batches. The relative frequency of individual shRNAmiRs in the cell population before and after reprogramming was assessed via sequencing. **A** Frequency distribution of shRNAmiRs in transduced parental cells and the 3 iPSC batches. **B** Enrichment of shRNAmiRs displayed as fold change of shRNAmiR frequencies in iPSC batches compared to transduced starting EC population. Red marked shRNAmiRs significantly (p < 0.001) derivate from sample median (one sample, wilcoxon signed rank test).

We observed 19.1 and 7.7 fold enrichment of one of the positive control shRNAmiRs ZFHX3.509 in 2 batches. Similarly, cells carrying shRNAmiRs SALL1.4587 and SALL1.110 were considerably enriched (5.7 and 6.4 fold), confirming our finding that actionable mutations in the SALL1 gene facilitate reprogramming. Also KLHL14.3195 appeared enriched in 2 batches, but the underlying mechanism remained unclear since literature research did not reveal any obvious connection of KLHL14 to pluripotency, cell cycle or apoptosis. In contrast, no positive selection (fold change < 2 in all batches) or enrichment in one batch only was detected for all other shRNAmiRs. Notably, in batch 3 one shRNAmiR SCEL.2743 took over almost the whole culture. However, we presume that this result rather represents a clonal event of insertional mutagenesis since enrichment is observed for one shRNAmiR in batch 3, only.

In summary, our results from EC-derived early passage iPSCs indicate that 98% of all iPSC small genetic variants are acceptable population polymorphisms that are passed over to iPSCs from founder cells. Among the remaining 2% of donor-specific variants we detected on average 2.7 (range 0-8) putatively actionable iPSC-specific mutations per clonal iPSC line. Furthermore, our analyses show that all analyzed true genetic variants detected in early passage iPSC lines pre-existed in the corresponding parental cell populations at different frequencies and indicate a high level of genetic mosaicism in human tissue. Our results also substantiate that there is no appreciable contribution of a hypothesized inherent mutagenicity of the reprogramming process to the mutational load in iPSCs.

Most importantly, we demonstrate for the first time the enrichment of SNVs and small INDELs from rare subpopulations of parental cells during reprogramming. Apparently, mutated clones outperform other clones during reprogramming and ultimately take over cultures of early reprogrammed cells. Our observations substantiate the finding of Shakiba et al. (Shakiba et al., 2019) by shedding light on the genetically encoded inequalities during the process of molecular reprogramming and by identifying potential driver genes affected by small-scale mutations. Among the group of enriched mutations, a substantial proportion affected genes involved in control of cell cycling, cell death and pluripotency. Although the numbers of variants were too low to reach statistical significance, putatively actionable mutations in oncogenes / tumor suppressor genes were apparently overrepresented among the group of enriched variants. For the tumor suppressor and pluripotency-specific gene SALL1, which was affected by one of the identified, enriched and putatively actionable donor-specific variants, we confirmed experimentally that downregulation can lead to clonal selection during reprogramming.

Although we did not observe a generally increased total number of SNVs and INDELs in iPSCs from elderly donors, our data show at least a trend for an increased number of putatively actionable variants in iPSCs from aged donors compared to neonatal cell sources, suggesting a lower biological quality of iPSCs from aged individuals (Lo Sardo et al., 2016; Ma et al., 2014).

## Limitations of Study

Our analysis was limited to 30 iPSC clones, all derived from ECs applying one reprogramming approach. Other cell sources and reprogramming techniques may result in different results. In any case, our findings indicate the requirement for further research to clarify the clinical risks accompanied with mutations that become enriched during reprogramming, such as malignant transformation of iPSC transplants. In particular, this includes the establishment of comprehensive databases of putatively actionable, enriched genetic variants in iPSCs, and their linking to databases in cancer research.

To generate comprehensive databases of genes that facilitate, when mutated, reprogramming and potentially pathogenic transformation, the investigation of a large number of iPSC lines will be necessary. Furthermore, it will be important to verify experimentally the effect of all genes affected by enriched mutations, as done exemplarily for SALL1, and to further investigate molecular mechanisms of enrichment. Lastly, reprogramming methods and applied culture conditions should be optimized to avoid or at least minimize the selection of cell clones that carry cancer and disease-causing mutations, which was also not focus of our study.

## Methods

### Isolation and culture of parental cells

ECs were chosen as donor cells, since these were available from different sources from neonatal (umbilical vein, hUVEC; cord blood, hCBEC) and aged (64-88 years, saphenous vein, hSVEC; peripheral blood, hPBEC) individuals. hUVECs and hSVECs were isolated from umbilical veins as well as hSVEC from saphenous veins using a standard enzymatic digestion protocol (Martin et al., 1998). hCBECs and hPBEC were isolated essentially as previously described (Haase et al., 2009). The ECs were cultured in EGM-2 medium (Lonza) for 4-5 passages, which equals to around 8-10 additional cell divisions after isolation, before reprogramming or subjecting to whole exome sequencing. Human material was collected after approval by the local Ethics Committee and following the donor’s written informed consent, or in the case of newborns, following parental consent.

### Virus Production

Plasmids pSIN-EF2-Lin-Pur, pSIN-EF2-Nanog-Pur, pSIN-EF2-Oct4-Pur, pSIN-EF2-Sox2-Pur (OSLN vectors) were purchased from Addgene. Viruses were produced, concentrated and titrated as described previously (Haase et al., 2009).

### Generation of hiPSCs

2×10^5^ ECs of passage 4-5 were reprogrammed by lentiviral transduction (multiplicity of infection, MOI 20) and characterized essentially as previously described (Haase et al., 2009). On day 6, the cells were transferred onto murine embryonic fibroblast (MEF) feeder layers and cultured with iPSC medium from day 7 onwards. For single cell cloning, individual colonies were manually transferred into separate wells and further cultured on MEFs. In total, 30 EC-derived iPSC clones were generated from 9 donors equating to a number of 3 iPSC clones per donor, with exception of donor D#37 for which 6 iPSC clones were generated (3 iPSC clones from each hSVEC and hPBEC).

### Genomic DNA extraction and whole exome sequencing

Genomic DNA (gDNA) was isolated from all 30 iPSC clones and EC parental cell population of 2 neonatal (D#2 hUVEC and D#3 hUVEC) and 2 aged (D#37 hSVEC and D#38 hSVEC) donors using Qiagen Blood DNA mini kits. Exomes were enriched using the commercially available Agilent SureSelectXT2 Human All Exon v4 (Agilent Technologies). 2µg of gDNA (> 3*10^5^ cells) of each sample were sheared and size selected to an average length of 200bp via AMPure XP bead purification. After adaptor ligation, the fragments were amplified for 8 cycles, purified and hybridized for 36h with an Exome capture library (Agilent, Santa Clara, USA; Human All Exon v4, consisting of biotinylated RNA probes). The DNA-biotinylated RNA hybrids were bound via streptavidin to magnetic beads, purified and amplified for 8 cycles with barcode containing primers. The final concentration, fragment distribution, and quality of the exome libraries were assessed on Qubit fluorimeter (Invitrogen) and Agilent Bioanalyzer. The sequencing was performed on a SOLiD 5500XL (Applied Biosystems, USA) with an Exact Chemistry Call (ECC) module. We sequenced 2 technical replicates with separate exome DNA amplification, barcode binding, emulsion PCR, bead enrichment and sequencing for each sample.

### Read alignment and variant calling

Reads (75bp, single-end) generated by the SOLiD 5500XL sequencer were aligned to the human reference genome hg19 using NovoalignCS (v1.04.02). Reads mapping equally well to more than one genomic location were discarded. The reads were trimmed during the alignment using the –H and –s 2 parameters of NovoalignCS. Reads with ≥ 4 nucleotides within the first 20bp or with ≥ 20% of all bases of Phred quality score < 10 (Ewing and Green, 1998) were discarded and PCR duplicates eliminated. Mapping quality and coverage were assessed employing Qualimap (v2.2.1). Variants were called in multisample format (coverage ≥ 10) grouping all 6 or 8 samples per donor (3 iPSC clones as replicates and, if so, 2 replicates of parental cell population) utilizing FreeBayes (v1.0.2).

### Variant validation and refinement

VCF files with FreeBayes results were imported into Galaxy (v17.05) (Afgan et al., 2018) instance of the RCU Genomics, Hannover Medical School, Germany for variant refinement. Complex variants were parsed running vcfAllelicPrimitives (v1.0.0_rc1.0) (Garrison 2015 (https://github.com/ekg/vcflib)) Galaxy built-in tool for simplification of variant refinement and annotation process.

The multisample variant calling allowed us to define variants with higher confidence also at a low coverage by applying a reverse variant refinement strategy. In detail, assuming inheritance from parental cell population the most variants would exist in all 3 iPSC clones. Therefore, a variant was regarded as being present in all iPSC clones unless its absence in one iPSC clone was defined by distinctively lower variant allele frequency (AF) in both replicates compared to the other samples of the same donor.

For developing our variant refinement strategy, we chose an empirical approach based on identification of true- and false-positive and -negative variants making use of variant phylogeny, meaning the dispersion of a variant in iPSC clones of one donor, and orthogonal validation sequencing (amplicon and Sanger sequencing). That means, the false-positive and -negative variants were identified by cross validation sequencing or by the presence or absence in one iPSC clone in which it would be expected to be or not to be according to the distribution of most other variants among the iPSC clones. This strategy was designed with the pivotal goal of returning variants shared by 2 iPSC clones with confidence. For a first variant validation round we applied no additional filter parameter. Polymorphisms were annotated utilizing Ensembl variant effect predictor (VEP) (v95) for human GRCh37 (assembly GRCh37.p13) based at European Bioinformatics Institute (EMBL-EBI) (McLaren 2016) with the GnomAD (gnomAD=170228) and 1000 Genome (dbSNP=151) modules and SnpSift Annotate (Cingolani et al., 2012) as Galaxy implemented tool with GnomAD (gnomad.exomes.r2.1) and dbSNP (149, reference GRCh37.p13). Polymorphisms defined by minor allele frequency (MAF) ≥ 0.01 in any population of GnomAD or 1000 Genome are termed in the following as “polymorphisms”. Moreover, a number of variants were found in more than one donor representing most likely either undescribed polymorphisms or sequencing error and barcode leakage and are hereafter termed “common variants”. We selected 84 variants (55 polymorphisms, 21 common variants and 8 donor-specific variants of which some showed low multisample quality value, sample coverage or alternative allele frequency (AF) (Table S1) for cross-validation by amplicon sequencing. All primers used for amplicon assays are listed in Table S7. For a more detailed description of the amplicon sequencing approach see below. The majority of polymorphisms, were confirmed as homo- or heterozygote variants in iPSC clones and parental cell populations. However, a number of variants of the group of uncertain and common variants, especially SNVs, were found to be false-positive (Table S1A). By applying a filter on multisample quality (≥ 400 and 530 for 6 and 8 sample batches, respectively) and DP (≥ 24 and 32) only around 1/5 of those false-positive variants could be eliminated. To exclude further false-positive variants, AF of variants was introduced as a filter criterion. For calculation of AF, multiallelic variants were excluded and a minimum sample DP was set. With DP6 per sample the allelic fractions of heterozygote variants followed roughly a binominal distribution with mean variant frequency of 0.45 (Fig. S1B) as expected (Genovese et al., 2014; Merkle et al., 2017). In the next step, first variants that were not in at least one iPSC clone detected with AF ≥ 0.3 were excluded as such proved demanding in the evaluation process as they often fell in difficult genomic regions and were error prone. Then different AFs thresholds (0.3, 0.2, 0.1 and adaptable) were tested in D#3 hUVEC iPSC clones as parameters for exclusion. Thereby, variants detected in mulitsample calling were assumed to be present in all iPSC clones of the donor unless AF in both replicates of an iPSC clones were below the set AF threshold. For all different threshold AFs the number of true- and false-positive and -negative variants was counted and error rates calculated. The assessment of true and false calls was based upon variant phylogeny (variants exist in iPSC clone 1 and 3 that originate apparently from same subpopulation, but not in iPSC clone 2), manual curation by differences in AF between samples, and supported by results of Sanger sequencing. While minimum AF 0.3 and AF 0.2 as threshold yield many false-negative variants, a threshold of AF 0.1 left false-positive variants. Error rate was lowest with adaptable AF (Fig. S1C), which considers differences between AFs of iPSC clones. In detail, a variant call in one clone was rejected if either i) AF in both replicates was < 0.1, or ii) AF in one replicate was < 0.15, undetected in other replicate, and AF > 0.4 in both replicates of both other clones, or iii) AF in both replicate < 0.15 and AF > 0.5 in both replicates of both other clones, or iv) AF in one replicate < 0.2, and undetected in other replicate, and AF > 0.5 in both replicates of both other clones, or v) AF in both replicates < 0.2 and AF > 0.6 in both replicates of both other clones.

After applying the new filter criteria, most false-positive variants were excluded (Table S1). However, a number of true-positive INDELs did not pass especially quality criterion (Table S1B). Although, here all analyzed INDELs were common polymorphisms or variants and not interesting for our further analysis, INDEL detection remains more demanding and less sensitive. Lastly, 10 variants of the passing 61 were not confirmed in iPSC and/or parental cells and likely represented sequencing errors. Those belonged, in general, to the group of common variants (Table S1B).

In the parental cell population, a variant was considered as not pre-existent if AF < 0.05 in both samples. The final workflow is displayed in Fig. S1A.

### Determination of variant allele fraction in parental cell population by amplicon sequencing

The error rate of around 1% of “Sequencing by Synthesis” NGS systems hinders the identification of rare genomic variants against a large background of non-mutated DNA sequences. Scientists have tried to overcome this problem mathematically: statistical methods have been applied to find significant imbalances in the pattern of sequencing errors and, thus, to identify and quantify these rare events. Unfortunately, this method is biased. If a statistically significant anomaly is found, this approach is robust and the analyzed variants can be considered verified. But the converse argument does not hold true: an insufficient significance is not proof for the absence of the respective variant. The smaller the mutated cell population is the higher the probability of missing this specific mutation event amid the vast statistical noise.

We developed an amplicon sequencing assay that combines amplification with high fidelity DNA polymerase and the highly accurate “Sequencing by Ligation” NGS technology to validate candidate variants in founder cell populations and to assess their allelic frequencies. PCR primers (Table S7) to amplify the regions spanning the variants were designed utilizing Primer3 (v0.4.0). In order to minimize the individual error rates for each sequenced base, we used the most accurate high fidelity DNA polymerase available for the initial amplification. 100 ng gDNA was used as template in the initial PCR amplification step, which corresponds to the genomes of > 1.6*10^4^ cells.

For INDEL (> 5bp) validation, PCR products of 1400-1600bp were generated by running 20 PCR cycles with a Q5 Hot Start High Fidelity DNA Polymerase (NEB GmbH, Frankfurt, Germany, error rates <10-7 per base). PCR products were purified using gel electrophoresis, extracted (QIAquick Gel Extraction Kit; Qiagen), and sheared on a Covaris sonifier according to manufacturer’s protocols to approximately 200bp fragments. INDEL fragment libraries were generated using the Fragment Library Core Kit (Life Technologies) as described by the manufacturer.

Similarly, PCR products, 60-120bp long, comprising SNV and INDELs ≤ 5bp were generated and purified. Fragment end repair of PCR products and adaptor ligation were performed with NEBNext Ultra II DNA Library Prep Kit for Illumina according to manufacturer. Subsequent steps of library preparation were performed using Fragment Library Core Kit.

Concentration, quality and fragment length distribution of generated libraries was assessed on Qubit and Agilent Bioanalyzer. The sequencing of the amplicons was performed on a SOLiD 5500XL (Applied Biosystems, USA) with an Exact Chemistry Call (ECC) module. For SNVs, the error rate and detection limit of amplicon sequencing assay was determined experimentally for each genomic location spanning the variants. In a first step, to exclude false sequences and reduce background noise, reads with the individual PCR primer sequence of the primer closer located to the SNV location were directly extracted from fastq files. Within the extracted reads, the number of reads with the SNV enclosed by a 15bp long sequence ranging from position −7bp to +7bp was counted as well as the number of reads with the reference sequence. To assess the average error also the numbers of reads that comprised one of the other both nucleotides (not SNV or reference nucleotide) were determined. Furthermore, two additional 15bp sequences, one directly upstream and one downstream of the SNV, were analyzed and numbers of reads for every non-reference nucleotide in the middle of the sequence counted. The mean error rate and detection limit were calculated for every SNV. Thus, the assay allows a deep insight into rare variants without the need for statistical extrapolation beyond the resolution of the sequencing technique. Frequencies of INDELs were determined by extracting and counting reads for both INDEL and reference embedded by a 20bp sequence, directly from fastq files. In some rare cases the determination of variant allele frequency of SNVs and INDELs in parental cells was performed on Illumina MiSeq platform. Therefore, PCR fragments of 1400-1600bp length were generated and sheared as described above, libraries were prepared via NEBNext Ultra II DNA Library Prep Kit for Illumina according to manufacturer, and then, due to higher error rate of the Illumina system, allele frequencies and errors were assed in a 11bp genomic context instead of 15bp as described above.

### Variant validation in iPSC clones

As iPSC clones are derived of individual founder cells, a variant is theoretically homo- or heterozygous. Therefore, variant validation via amplicon sequencing in iPSC clones was performed with the same technical approach as described above for determination of variant fraction in parental cell population but the result interpretation was simplified. Only reads with the SNV enclosed by a 15bp sequence and the reference sequence were quantified, but no error rate was estimated. Validation of INDELs was performed as described above.

For validation of variants via Sanger sequencing, PCR products were prepared as for amplicon sequencing. Purified PCR products were prepared according to company’s requirement and sent together with the PCR primers to Microsynth Seqlab (Göttingen, Germany). Results were visualized via Chromas software (v1.45).

### Variant annotation and prediction of functional consequences

Variant annotation was performed using SnpEff (hg19) (v4.3) (Cingolani et al., 2012) as Galaxy implemented tool and the web interface of Ensembl VEP (v95) (assembly GRCh37.p13) based at European Bioinformatics Institute (EMBL-EBI) (McLaren et al., 2016). The functional consequence of a GAVs was classified by a consensus based on the in silico predictions of Condel, FATHMM, CADD (Ensembl modules) and SnpEff impact. If a variant had a harmful designation by SnpEff (high impact) or at least by two of the algorithms (Condel=deleterious, FATHMM=damaging, CADD phred > 17), provided prediction tools returned a result, it was considered as putatively actionable.

The number of all nucleotide change events in the 3 collective of GAVs found in 1, 2, or 3 iPSC clones of all donors were determined and mutational spectra compared to Signatures of Mutational Processes in Human Cancer of COSMIC Catalogue of Somatic Mutations in Cancer hosted by Wellcome Trust Sanger Institute.

GO process annotation of genes was obtained from GO biological process (C5BP) collection of Molecular Signatures Database (MSigDB) (Liberzon et al., 2015) maintained by Broad Institute. In case no GO process was curated, IntAct Molecular Interaction Database hosted by EMBL-EBI, BioGRID Biological General Repository for Interaction Datasets (Oughtred et al., 2019) and UniProt (UniProt, 2019) were consulted for interacting proteins and their GO processes. To estimate the oncogenic potential of each variant, we computed overlap of affected genes with MSigDB Collection of Computational gene sets (C4) and Oncogenic signatures (C6) as well as Jensen DISEASE database (Pletscher-Frankild et al., 2015). Furthermore, OncoKB database (Chakravarty et al., 2017) as well as the COSMIC Cancer gene census (Sondka et al., 2018) were utilized for matching of curated onco- and tumor suppressor genes.

### Analysis of clonal enrichment of shRNAmiRs during reprogramming

Lentiviral vector construction of pLKO5d.SFFV.eGFP.miR30N.WPRE vector was described previously (Adams et al., 2017). Within the miR-N cloning cassette the SFFV promotor was replaced by a short CAG promotor. shRNAmiR design, cloning and library construction, packaging into lentivirus and test via reporter assay were performed as previously described (Adams et al., 2017; Fellmann et al., 2013; Schwarzer et al., 2017). In short, for each target gene 3 shRNAmiRs were designed (Table S6). The 67mer oligonucleotides encoding the shRNAmiR construct including passenger (22bp), loop (19bp), guide (22bp) and overhang (4bp) were purchased form Integrated DNA Technologies (IDT) and cloned into the pLKO5d.CAGs.eGFP.miR30N.WPRE backbone. The shRNAmiR constructs were then mixed equimolarly to form shRNAmiR libraries and virus containing the library was produced and tested. D#2 hUVEC cells were transduced with shRNAmiR library with MOI 2, cultured for 3 passages and then reprogrammed with monocistronic lentiviral Thomson factors as described above. 3 technical replicates (batches of iPSCs) were derived from the same transduced parental cell starting population with 4*10^5^ transduced cells in each batch. 57, 53, and 64 iPSC colonies arose in the 3 batches, respectively, which equals to around 0.015% reprogramming efficiency. Reprogrammed cells were cultured for 3 passages and gDNA isolated (see above) form the entirety of reprogrammed cells (without sampling of cells during passaging).

Composition of the shRNAmiR library in the 3 iPSC batches compared to the transduced parental cell population was analyzed by sequencing. For this shRNAmiR sequencing, PCR product of 157bp, spanning the shRNAmiR sequence and parts of the 3’ and 5’ miR30A flank were generated by running 25 cycles with Q5 Hot Start High Fidelity DNA Polymerase (forward primer: GTTAACCCAACAGAAGGCTAAAG, reverse primer: TAATTGCTCCTAAAGTAGCCCCTTG; Annealing temperature 63°C). This PCR reaction amplified not only the shRNAmiR sequences but also the endogenous miRN30A of the cells. Sequencing library preparation was conducted on PCR fragments with NEBNext Ultra II DNA Library Prep Kit for Illumina according to manufacturer’s instruction. The quality of the fragment library was assessed on Qubit fluorimeter and Agilent Bioanalyzer. The sequencing was performed on Illumina MiSeq. Assessment of content of each individual shRNAmiR in iPSC batches and in transduced parental cells was performed by counting sequencing reads. For this, raw reads were filtered for 12bp of PCR primer sequence allowing 1 mismatch, and then endogenous miR30A reads were excluded (filtering for 12bp accepting 1 mismatch). Finally, the numbers of reads for each shRNAmiR were counted by filtering for a 15bp sequence of the guide shRNAmiR allowing two mismatches within this 15bp sequence.

### Statistical analyses

GraphPad Prism (v6.07) or RStudio (v1.1.463) software were utilized for graph visualization and data statistics. Data are given as mean ± SD. Sample distribution was assessed employing D’Agostino and Pearson omnibus normality test. Unpaired two-tailed t-tests were applied to compare number of variants found per iPSC clone between age groups. Nonparametric two-tailed Mann Whitney test was performed to compare number of putatively actionable variants found per iPSC clone between age groups.

For assessment of error rate of the amplicon sequencing assay, the mean value of incorrectly recorded nucleotides (n = 8) was calculated for every SNV. Distribution of errors for each SNV was evaluated via D’Agostino and Pearson omnibus normality test or, in rare cases, Shapiro-Wilk normality test if the first one was not applicable. In case given samples followed a normal distribution, a one sample one-tailed t-test was applied to compare read count of SNV as hypothetical value against background of incorrectly annotated bases. Otherwise, Wilcoxon one-tailed signed-rank test was employed. Null hypothesis which propose that read number of SNV lies within local error range was rejected with p value 0.1.

### Code availability

Variant refinement was performed on the internal Galaxy (v17.05) (Afgan et al., 2018) instance of the RCU Genomics, Hannover Medical School, Germany. All workflows that were used to process variants are available as supplemental file (Data S2). The utilized script for processing and analysis of shRNAmiR sequencing data is available in Data S3.

### Data availability

All data underlying the study are available on request and are currently deposited on servers of Hannover Medical School, Germany.

## Acknowledgements

The authors are grateful to J. Gorenaja, K. Menge, M. Sievert, M. Tauscher, U. Opel and, S. Mielke for their technical support, to P. Chouvarine for initial support regarding bioinformatics, to N. McGuinness for a critical reading of the manuscript, and to T. Scheper for providing bFGF. Some of human cord blood samples were kindly provided by Vita 34, Leipzig. Many thanks to C. Hess for the isolation of Vita34 CBECs and to P. Stiefel for several HSVEC isolations. We are also thankful to S. Rojas, T. Goecke and other surgical colleagues, who provided human tissue samples. We thank J. Thomson for providing the reprogramming plasmids via Addgene and D. Trono for providing the lentiviral packaging and transfer plasmids psPAX2 and pMD2.G.

## Author contributions

M.K. and K.O. performed experiments, collected and analyzed data and contributed to writing the manuscript. C.D. and L.W. performed WES, analyzed and collected bioinformatics data and contributed to writing the manuscript. F.A., A.Schw. and A.Scha. designed the shRNAmiR library. A.H., Sy.M. and S.W. contributed to the conduction of experiments. Sa.M. and M.D. provided technical assistance in cell culture, fragment library construction and WES. U.M. designed and coordinated the study and wrote the manuscript.

## Supplemental information

**Fig. S1:**
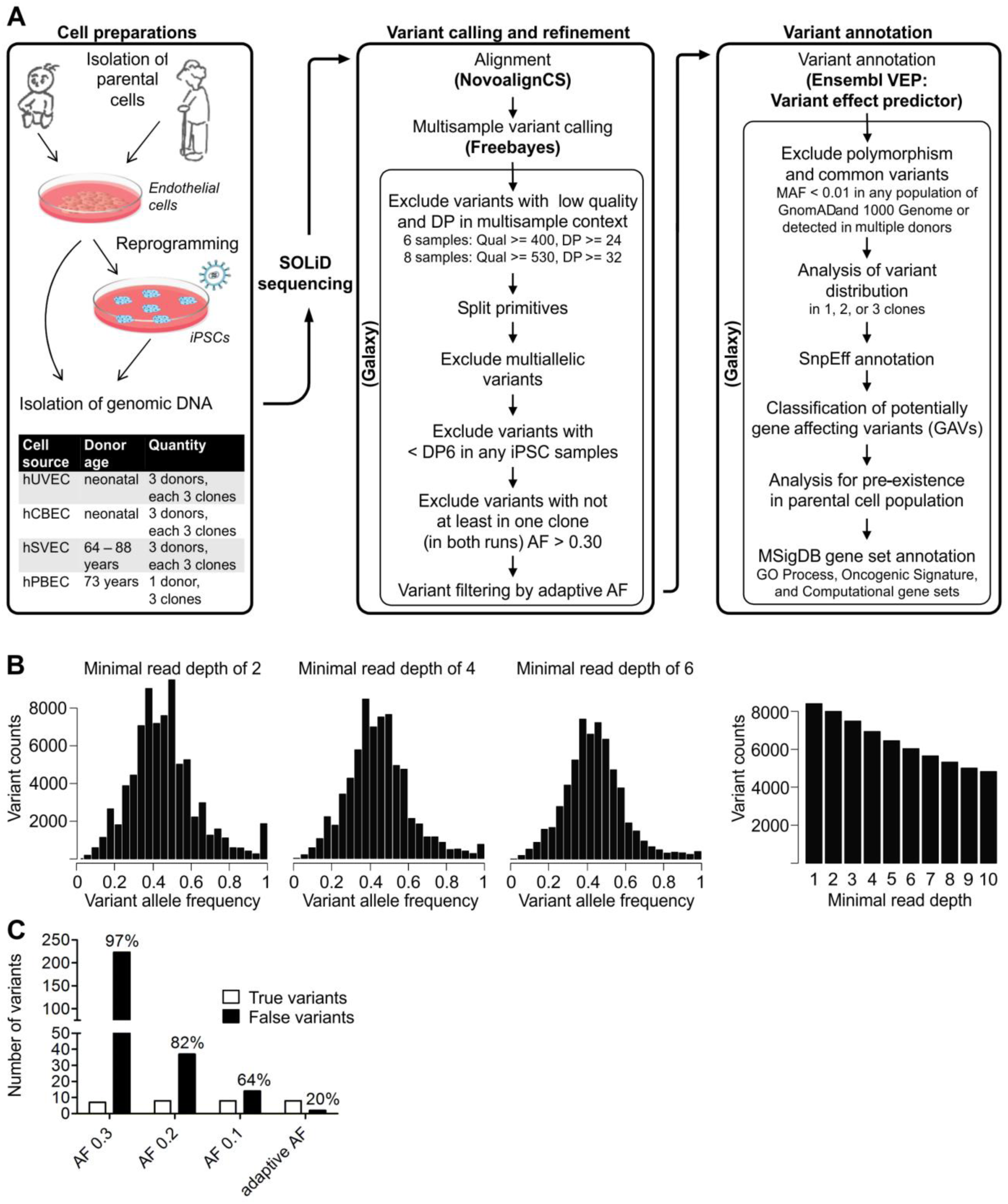
Establishment of variant refinement strategy and final workflow. **A** Workflow of whole exome sequencing (WES) performance and analysis including cell preparation, variant calling and refinement process, and variant annotation. Exomes from 30 early passage iPSC clones of neonatal and aged donors were sequenced twice as technical replicates. Similarly, exomes of corresponding parental cell populations of 2 of the neonatal and 2 of the aged donors were sequenced. Subsequent to variant calling and filtering by multisample quality value and coverage, further variant refinement based on sample coverage (DP) and variant allele frequency (AF) threshold was carried out. **B** Exemplarily for D#3 hUVEC C1, the AF distribution of all heterozygote variants is depicted after applying minimum sample coverage of DP1-DP10 (DP2, DP4, and DP6 displayed) as filters. With DP6 per sample the allelic fractions of heterozygote variants followed roughly a binominal distribution with mean variant frequency of 0.45 and was chosen as adequate filter parameter value for further processing. **C** Number and ratio of true variants (true-positive) and false variants (false-positive and negative) sheared by 2 iPSC clones, exemplarily, of donor D#3 hUVEC generated by filtering with different AF thresholds. True and false calls were manual curation based upon variant phylogeny and by differences in AF between samples, and supported by cross validation via Sanger sequencing. While minimum AF 0.3 and AF 0.2 as threshold yield many false-negative variants, a threshold of AF 0.1 left false-positive variants. Adaptive AF filter preserved all true-positive variants while reducing false calls to an acceptable number. Abbreviations: DP, depth (of coverage) AF, allele frequency; MAF, minor allele frequency; GO, gene ontology.

**Fig. S2:**
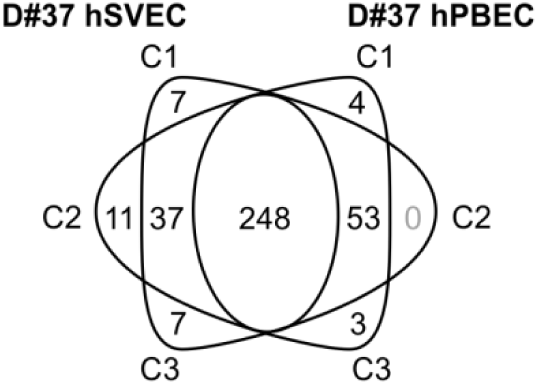
Tissue distribution of GAVs in donor D#37. Endothelial cells (ECs) were isolated from saphenous vein (hSVEC) and peripheral blood (hPBEC) of donor D#37. 3 iPSC clones from each, hSVEC and hPBEC, were generated and subjected to whole exome sequencing. Venn diagram showing for donor D#37 number of unique and shared donor-specific, potentially gene affecting variants (GAVs) (located in coding and non-coding transcript region, UTRs, and splice regions).

**Table S3:** **Dispersion of variants in iPSC clones.** 30 iPSC clones were generated from in total 10 neonatal and aged cell source and subjected at passage 7-10 to whole exome sequencing. Number and percentage of donor-specific variants and potentially gene affecting variants (GAVs) (without intergenic and intron variants) in 1, 2, and 3 iPSC clones were determined.

| Donor |  | Variants |  | GAVs |  |
| --- | --- | --- | --- | --- | --- |
|  |  | Total number | Percentage | Total number | Percentage |
| **iPSCs - Neonatal Donors** |  |  |  |  |  |
| hUVECiPSC D#1 | 1 clone | 13 | 2.70 | 11 | 4.28 |
|  | 2 clones | 1 | 0.21 | 1 | 0.39 |
|  | 3 clones | 468 | 97.10 | 245 | 95.33 |
| hUVECiPSC D#2 | 1 clone | 14 | 2.72 | 11 | 3.69 |
|  | 2 clones | 0 | 0.00 | 0 | 0.00 |
|  | 3 clones | 500 | 97.28 | 287 | 96.31 |
| hUVECiPSC D#3 | 1 clone | 12 | 1.57 | 9 | 2.41 |
|  | 2 clones | 8 | 1.05 | 6 | 1.61 |
|  | 3 clones | 742 | 97.38 | 358 | 95.98 |
| hCBECiPSC D#22 | 1 clone | 18 | 2.79 | 14 | 4.59 |
|  | 2 clones | 3 | 0.47 | 0 | 0.00 |
|  | 3 clones | 624 | 96.74 | 291 | 95.41 |
| hCBECiPSC D#23 | 1 clone | 11 | 2.04 | 6 | 1.97 |
|  | 2 clones | 1 | 0.19 | 1 | 0.33 |
|  | 3 clones | 527 | 97.77 | 298 | 97.70 |
| hCBECiPSC D#25 | 1 clone | 4 | 0.72 | 4 | 1.38 |
|  | 2 clones | 7 | 1.25 | 5 | 1.72 |
|  | 3 clones | 548 | 98.03 | 281 | 96.90 |
| Mean | 1 clone | 12 | 2.09 | 9 | 3.05 |
|  | 2 clones | 3 | 0.53 | 2 | 0.67 |
|  | 3 clones | 568 | 97.38 | 293 | 96.27 |
| **iPSCs - Aged Donors** |  |  |  |  |  |
| hSVECiPSC D#31<br>(64 years) | 1 clone | 15 | 3.23 | 10 | 3.88 |
|  | 2 clones | 9 | 1.94 | 5 | 1.94 |
|  | 3 clones | 441 | 94.84 | 243 | 94.19 |
| hSVECiPSC D#37<br>(73 years) | 1 clone | 41 | 6.54 | 25 | 8.06 |
|  | 2 clones | 1 | 0.16 | 0 | 0.00 |
|  | 3 clones | 585 | 93.30 | 285 | 91.94 |
| hSVECiPSC D#38<br>(88 years) | 1 clone | 27 | 2.85 | 19 | 6.17 |
|  | 2 clones | 23 | 2.43 | 13 | 4.22 |
|  | 3 clones | 896 | 94.71 | 276 | 89.61 |
| hPBECiPSC D#37<br>(73 years) | 1 clone | 7 | 1.13 | 7 | 2.27 |
|  | 2 clones | 1 | 0.16 | 0 | 0.00 |
|  | 3 clones | 612 | 98.71 | 301 | 97.73 |
| Mean | 1 clone | 23 | 3.44 | 15 | 5.10 |
|  | 2 clones | 9 | 1.17 | 5 | 1.54 |
|  | 3 clones | 634 | 95.39 | 276 | 93.36 |
| **iPSCs - All Donors** |  |  |  |  |  |
| Mean | 1 clone | 16 | 2.63 | 12 | 3.87 |
|  | 2 clones | 5 | 0.78 | 3 | 1.02 |
|  | 3 clones | 594 | 96.59 | 287 | 95.11 |

**Table S4:** **Pre-existence of variants in parental cell population.** Whole exome sequencing (WES) of endothelial cell population of 4 donors was performed to investigate the origin of donor-specific, potentially gene affecting variants (GAVs) (located in coding and non-coding transcript region, UTRs, and splice regions) that had been detected in 1, 2, or all 3 iPSC clones per donor by WES. Number and percentage of pre-existent and via WES undetected variant in parental cell population was determined. Thereby, allele frequency of 0.05 in parental cell population was the detection limit.

| Donor |  | Total number | Pre-existent in parental cell population | Pre-existent in parental cell population (%) | Undetected in parental cell population |
| --- | --- | --- | --- | --- | --- |
| **iPSCs - Neonatal Donors** |  |  |  |  |  |
| hUVECiPSC D#2 | 1 clone | 11 | 0 | 0 | 11 |
|  | 2 clones | 0 | 0 | 0 | 0 |
|  | 3 clones | 286 | 286 | 100.00 | 0 |
| hUVECiPSC D#3 | 1 clone | 9 | 0 | 0 | 9 |
|  | 2 clones | 6 | 0 | 0 | 6 |
|  | 3 clones | 358 | 356 | 99.44 | 2 |
| Mean | 1 clone | 10 | 0 | 0 | 10 |
|  | 2 clones | 3 | 0 | 0 | 3 |
|  | 3 clones | 322 | 321 | 99.72 | 1 |
| **iPSCs - Aged Donors** |  |  |  |  |  |
| hSVECiPSC D#37 (73 years) | 1 clone | 25 | 0 | 0 | 25 |
|  | 2 clones | 0 | 0 | 0 | 0 |
|  | 3 clones | 285 | 285 | 100 | 0 |
| hSVECiPSC D#38 (88 years) | 1 clone | 19 | 0 | 0 | 19 |
|  | 2 clones | 13 | 0 | 0 | 13 |
|  | 3 clones | 276 | 270 | 97.83 | 6 |
| Mean | 1 clone | 22 | 0 | 0 | 22 |
|  | 2 clones | 7 | 0 | 0 | 7 |
|  | 3 clones | 281 | 278 | 98.91 | 3 |
| **iPSCs - All Donors** |  |  |  |  |  |
| Mean | 1 clone | 18 | 0 | 0 | 10 |
|  | 2 clones | 4 | 0 | 0 | 5 |
|  | 3 clones | 282 | 280 | 99.28 | 2 |

## External Data S1: Excel file with Table S1, S2, S5, S6, and S7

**Table S1A: Cross-validation of uncertain variants that failed criteria of the final variant refinement strategy.** 30 iPSC clones of neonatal and aged donors were subjected to whole exome sequencing twice as technical replicates. Following variant calling, polymorphisms were annotated. While (category A) variants defined as polymorphisms are described as such in any population of GnomAD and 1000 Genome with minor allele frequency (MAF) ≥ 0.01, common (category B) variants are such detected in more than 1 donor and might be as yet unknown polymorphisms or sequencing artefacts. Orthogonal sequencing for cross-validation was performed for determination of both, optimal variant refinement strategy and dimension of false variant retention and true variant elimination. Exemplarily, genomic regions of 23 uncertain variants (9 polymorphisms, 6 common variants, and 8 donor-specific variants) that failed filtering by criteria of our final variant refinement strategy were PCR amplified from gDNA of iPSC clones and validated via amplicon (*) or Sanger sequencing. The column “Failing criteria in variant validation” expresses whether a variant was eliminated due to low multisample quality value (Qual), low depth of coverage (DP < 6), multiallelic calls (MC), or allele frequency < 0.3 in at least 1 replicate in all 3 iPSC clones (AF). Cross-validation demonstrated that the half of these uncertain variants could not be confirmed (upper part). The other half comprised primarily true INDELs (polymorphisms) that were eliminated due to low mapping quality (lower part).Abbreviations: DP, depth (of coverage); AF, allele frequency; MC, multiallelic calls, amb, ambiguous.

**Table S1B: Cross-validation of variants that passed criteria of the final variant refinement strategy and their allele frequency in parental cells.** 30 iPSC clones of neonatal and aged donors were subjected to whole exome sequencing twice as technical replicates. Following variant calling, polymorphisms were annotated. While (category A) variants defined as polymorphisms are described as such in any population of GnomAD and 1000 Genome with minor allele frequency (MAF) ≥ 0.01, common (category B) variants are such detected in more than 1 donor and might be as yet unknown polymorphisms or sequencing artefacts. Orthogonal sequencing for cross-validation was performed for determination of both, optimal variant refinement strategy and dimension of false variant retention and true variant elimination. Genomic regions of 61 variants (46 polymorphisms, and 15 common variants were PCR amplified from gDNA of parental cell population and analysed via amplicon sequencing. As in parental cell population variants might exist only in subpopulations and, consequently, at low allelic fraction, pre-existence of variants was evaluated closely (with a p-value of 0.1) taking local error rates into account. The existence of almost all polymorphisms could be confirmed as homo- or heterozygote variant within the corresponding parental cell populations by amplicon sequencing (upper part). Existence of a representative choice of variants in iPSC clones was cross-validated via amplicon (*) or Sanger sequencing. This orthologous sequencing for cross-validation was performed for determination of both, optimal variant refinement strategy and dimension of false variant retention and true variant elimination. Cross-validation demonstrated that only a small number of false variants calls pass final filter strategy (lower part). Those false variants primarily belong to the group of common variants. () Pre-existence in parental cells not confirmed with statistical confidence (p-value > 0.1).

**Table S2A: Determination of variant allele frequencies of SNVs in parental cell populations.** Amplicon sequencing of potentially gene affecting variant (GAV) (located in coding and non-coding transcript region, UTRs, and splice regions) spanning regions in parental cell populations was performed. GAVs belong to polymorphisms with minor allele frequency (MAF) ≥ 0.01 in any population of GnomAD and 1000 Genome (Cat. A), common variants (detected in more than 1 donor; Cat. B), or donor-specific (Cat. C) variants and were found in 1, 2, or all 3 iPSC clones. The table list SNVs and their allele frequencies in iPSC clones and parental cell population determined by whole exome sequencing. If conducted, confirmation of SNVs in iPSC clones was performed by amplicon or Sanger sequencing. Amplicon sequencing results for SNVs are displayed as the number of reads per respective nucleotide embedded in a 15bp sequence. The reference allele reads are given in bold, and the variant genotype reads in red. Read counts for error rate calculation, average local error rates, information to applied statistical test and results, as well as local detection limits are presented.

**Table S2B: Determination of variant allele frequencies of INDELs in parental cell populations.** Amplicon sequencing of potentially gene affecting variant (GAV) (located in coding and non-coding transcript region, UTRs, and splice regions) spanning regions in parental cell populations was performed. GAVs belong to polymorphisms with minor allele frequency (MAF) ≥ 0.01 in any population of GnomAD and 1000 Genome (Cat. A), common variants (detected in more than 1 donor; Cat. B), or donor-specific (Cat. C) variants and were found in 1, 2, or all 3 iPSC clones. The table list INDELs and their allele frequencies in iPSC clones and parental cell population determined by whole exome sequencing. If conducted, confirmation of INDELs in iPSC clones was performed by amplicon or Sanger sequencing. Evaluation of amplicon sequencing results for INDELs comprised determination of reads for reference and variant embedded by a 20bp sequence.

**Table S5A: List of enriched donor-specific GAVs detected in 2 or 3 out of 3 iPSC clones per donor including information regarding predicted effects and affected genes.** The table lists all potentially gene affecting variants (GVAs), that were found in 3 or 2 out of 3 iPSC clones but were not detected in parental cell population by WES (upper part), or were found in 2 out of 3 iPSC clones, but were not analyzed in the parental cell population via WES (lower part). In contrast to Table 2 this includes putatively actionable and neutral variants. Variant allele frequencies in iPSC clones were determined by whole exome sequencing (WES). Confirmation of variants in iPSC clones was performed by amplicon or Sanger sequencing. WES of parental cells of 4/10 donors (D#2 hUVEC, D#3 hUVEC, D#37 hSVEC and D#38 hSVEC) revealed that variants detectable in less than 3 iPSC clones of the individual donors were always undetectable in the parental cells. Moreover, in some cases, also pre-existence of variants detected in all 3 iPSC of a donor could not be confirmed by WES. A more sensitive amplicon sequencing of GAV spanning regions for precise determination of allelic frequencies of GAVs in corresponding parental cell population, which was performed for a representative choice of GAVs, however, demonstrated the presence of all tested GAVs in the parental cells at low frequency. Pre-existence of variants in parental cell population was evaluated (with a p-value 0.1) taking local error rates into account. Since entirely all variants that were detected in 1 or 2 out of 3 iPSC generated from the above 4 donors were demonstrated to be not detectable by WES but were detectable by amplicon sequencing, variants detected in 2 out of 3 iPSC clones derived from the remaining 6 donors D#1 hUVEC, D#22 hCBEC, D#23 hCBEC, D#25 hCBEC, D#31 hSVEC and D#37 hPBEC were assumed to follow the same pattern. The functional consequence of donor-specific GAVs was classified by a consensus of the in silico prediction of Condel, FATHMM, CADD and SnpEff impact. Variants were classified as putatively actionable (marked in red under “Effect prediction”), if SnpEff returned a high impact prediction or at least two of the other tools a harmful designation, or otherwise as neutral (marked in blue) or ambiguous (marked in purple). GO process annotations and notion of cancer-relation of affected genes were retrieved from MSigDB Collections of GO biological process, Computational gene sets and Oncogenic signatures as well as Jensen DISEASE database. Curated onco- or tumor suppressor genes (annotation retrieved from OncoKB or COSMIC Cancer gene census) are shaded in red. Variants that are presumed to influence reprogramming since they affect genes with function in control of cell cycling, cell death or pluripotency are depicted in blue font. () Pre-existence in parental cells likely but not confirmed with statistical confidence (p-value > 0.1).

**Table S5B: List of donor-specific GAVs detected in 1 out of 3 iPSC clones per donor including information regarding predicted effects and affected genes.** The table lists all GAVs that were found in 1 out of 3 iPSC clones but were not detected (upper part) or not analyzed (lower part) in parental cell population via WES. Variant allele frequencies in iPSC clones were determined by whole exome sequencing (WES). Confirmation of variants in iPSC clones was performed by amplicon or Sanger sequencing. WES of parental cells of 4/10 donors (D#2 hUVEC, D#3 hUVEC, D#37 hSVEC and D#38 hSVEC) revealed that variants detectable in 1 iPSC clone of the individual donors were always undetectable in the parental cells. A more sensitive amplicon sequencing of GAV spanning regions for precise determination of allelic frequencies of GAVs in corresponding parental cell population, which was performed for a representative choice of GAVs, however, demonstrated the presence of all tested GAVs in the parental cells at low frequency. Pre-existence of variants in parental cell population was evaluated (with a p-value 0.1) taking local error rates into account. The functional consequence of donor-specific GAVs was classified by a consensus of the in silico prediction of Condel, FATHMM, CADD and SnpEff impact. Variants were classified as putatively actionable (marked in red under “Effect prediction”), if SnpEff returned a high impact prediction or at least two of the other tools a harmful designation, or otherwise as neutral (marked in blue) or ambiguous (marked in purple). GO process annotations and notion of cancer-relation of affected genes were retrieved from MSigDB Collections of GO biological process, Computational gene sets and Oncogenic signatures as well as Jensen DISEASE database. Curated onco- or tumor suppressor genes (annotation retrieved from OncoKB or COSMIC Cancer gene census) are shaded in red. Variants that are presumed to influence reprogramming since they affect genes with function in control of cell cycling, cell death or pluripotency are depicted in blue font.

**Table S6: Relative frequency of shRNAmiRs before and after reprogramming.** ECs were transduced with a library containing shRNAmiRs against a choice of 16 genes (3 shRNAmiRs per gene). Transduced cells were reprogrammed in 3 independent batches and the relative frequency of shRNAmiRs in the EC population and the 3 iPSC batches was determined via sequencing. The table displays shRNAmiR sequences, the read counts for each shRNAmiR and the calculated fold change of shRNAmiRs in iPSC batches compared to EC population. shRNAmiRs against genes that were affected by a putatively actionable enriched GAVs are highlighted in bold typeface and blue, such against neutral enriched GAVs are shown in bold.

**Table S7: Primer list.** Primer sequences for amplicon and Sanger sequencing of every analyzed variant are provided including product length and annealing temperature.

External Data S2: Galaxy workflows.

External Data S3: Script for shRNAmiR analysis.

